# Dual control of host actin polymerization by a *Legionella* effector pair

**DOI:** 10.1101/2023.05.15.540800

**Authors:** M. Pillon, C. Michard, N. Baïlo, J. Bougnon, K. Picq, O. Dubois, C. Andrea, L. Attaiech, V. Daubin, S. Jarraud, E. Kay, P. Doublet

**Affiliations:** CIRI, Centre International de Recherche en Infectiologie, (Team : Legionella pathogenesis), Univ Lyon, Inserm, U1111, Université Claude Bernard Lyon 1, CNRS, UMR5308, ENS de Lyon, F-69007, Lyon, France; CIRI, Centre International de Recherche en Infectiologie, (Team : HORIGENE), Univ Lyon, Inserm, U1111, Université Claude Bernard Lyon 1, CNRS, UMR5308, ENS de Lyon, F-69007, Lyon, France; Laboratoire de Biométrie et Biologie Evolutive, Université de Lyon, 69000 Lyon, France Centre National de la Recherche Scientifique, Unité Mixte de Recherche 5558, Université Lyon 1, 69622 Villeurbanne, France

**Author notes:** Corresponding author: Pr. Doublet Patricia; e.mail.

**Keywords:** *Legionella pneumophila*, secretion effectors, effector-effector functional interaction, host actin cytoskeleton, endosomal pathway, LegK2, VipA, effector repertoire

## Abstract

Host actin cytoskeleton is often targeted by pathogenic bacteria through the secretion of effectors. *Legionella pneumophila* virulence relies on the injection of the largest known arsenal of bacterial proteins, over 300 Dot/Icm Type 4 Secretion System effectors, into the host cytosol. Here we define the functional interactions between VipA and LegK2, two effectors with antagonistic activities towards actin polymerization that have been proposed to interfere with the endosomal pathway. We confirmed the prominent role of LegK2 effector in *Legionella* infection, as the deletion of *legK2* results in defects in the inhibition of actin polymerization at the *Legionella* Containing Vacuole, as well as in endosomal escape of bacteria and subsequent intracellular replication. More importantly, we observed the restoration of the *ΔlegK2* mutant defects, upon deletion of *vipA* gene, making LegK2/VipA the first example of effector-effector suppression pair that targets the actin cytoskeleton and whose functional interaction impacts *L. pneumophila* virulence. We demonstrated that LegK2 and VipA do not modulate each other’s activity in a ‘metaeffector’ relationship. Instead, the antagonistic activities of the LegK2/VipA effector pair would target different substrates, Arp2/3 for LegK2 and G-actin for VipA, to temporally control actin polymerization at the LCV and interfere with phagosome maturation and endosome recycling, thus contributing to the intracellular life cycle of the bacterium. Strikingly, the functional interaction between LegK2 and VipA is consolidated by an evolutionary history that has refined the best effector repertoire for the benefit of *L. pneumophila* virulence.

## INTRODUCTION

Actin cytoskeleton, including actin itself (globular, filamentous and its polymerization) as well as accessory proteins such as myosins, is a preferential target for pathogenic bacteria^1–3^. It is a complex and dynamic network that shapes the cell and plays a key role in numerous cellular processes such as cell migration, adhesion, internalization and intracellular trafficking. Intracellular bacteria evolved effective strategies that take advantage of actin polymerization in order (i) to gain entry into epithelial cells^4–7^, (ii) to promote their movement within the host cytosol, thus contributing to their evasion from autophagy and their propagation to neighboring cells^8,9^, and (iii) to stabilize their replicative vacuole in epithelial cells^10,11^. Actin polymerization inhibition or actin degradation strategies have also been described to contribute to bacterial evasion from phagocytosis^12^ and to cell cytotoxicity and therefore pathogenesis development^13^.

*Legionella pneumophila*, the etiological agent of the severe pneumonia legionellosis, is a typical example of intracellular pathogen that has evolved several strategies to take advantage of the host actin cytoskeleton. This sophisticated relationship allows the pathogen to achieve intracellular replication within phagocytic cells such as amoebae in aquatic environment or alveolar macrophages in its accidental human host. Intracellular replication occurs in a rough endoplasmic reticulum-like compartment called *Legionella* Containing Vacuole (LCV) that evades endocytic maturation. The Type 4 Secretion System (T4SS) Dot/Icm is crucial for *Legionella* intracellular replication, through the secretion of over 300 effectors into the host cell cytoplasm. To date, five Dot/Icm effectors have been shown to interfere with actin polymerization, activating it for VipA, inhibiting it for the others. Specifically, VipA is an actin nucleator, which localizes to early endosomes during infection and promotes actin polymerization *in vitro*^14^; Ceg14 co-sediments with F-actin and inhibits actin polymerization by an unknown mechanism^15^; LegK2 is a Ser/Thr kinase that phosphorylates ArpC1b and Arp3, two subunits of the Arp2/3 complex, thus inhibiting actin polymerization on the LCV and contributing to the bacterial evasion from endosomal degradation^16^; RavK is a metalloprotease that cleaves actin, generating a fragment with a diminished capacity to form actin filaments^17^; and WipA is a phosphotyrosine that disrupts F-actin polymerization by hijacking phosphotyrosine signaling^18^.

We sought to establish the functional interactions between VipA and LegK2, as both effectors exhibit antagonistic activities towards actin polymerization and both have been proposed to interfere with the endosomal pathway^14,16^. We constructed and characterized simple and double mutants with in-frame deletion of the genes encoding these effectors. We confirmed the role of LegK2 in the inhibition of actin polymerization on the LCV. More importantly, we observed the restoration of the *ΔlegK2* mutant defect in this step, upon deletion of *vipA*, thus identifying the first *Legionella* effector-effector suppression pair targeting the host cell actin cytoskeleton. By doing so, we proposed for the first time a role for VipA in the infectious cycle of *L. pneumophila*. Strikingly, the compensation of the *legK2* phenotype by *vipA* deletion was also shown for bacterial escape from endosomal pathway as well as for intracellular replication, thus making LegK2/VipA the first example of effector-effector suppression pair whose functional interaction impacts *L. pneumophila* virulence. We demonstrated that LegK2 and VipA differ from other effector-effector suppression pairs identified by targeted functional studies or systematic screens^19^, in that these effectors do not exhibit antagonistic enzymatic activities against a common substrate, and do not modulate the other’s activity in a metaeffector-effector relationship. Rather, they target different components of the actin cytoskeleton, to contribute to the intracellular life cycle of the bacteria. Finally, combined with the phylogenetic study of the genes encoding LegK2 and VipA, this work shows that the functional interaction between two Dot/Icm effectors results from an evolutionary history that has refined the best effector repertoire for the benefit of *L. pneumophila* virulence.

## MATERIALS AND METHODS

### Cell lines & Bacterial strains

Bacterial strains, cells lines used in this study are summarized in Supplementary Data S1. *Legionella pneumophila* strains were grown at 37°C either in BCYE (buffered charcoal yeast extract) agar or in AYE (ACES ([*N*-(2-acetamido)-2-aminoethanesulfonic acid] Yeast Extract) liquid medium (12 g/L yeast extract (Difco), 10 g/L ACES (Roth), 0.4 g/L L-cystein HCl (Euromedex), 0.25 g/L, pH adjusted to 6.9 with KOH 1N). Each medium was supplemented with 5 μg/ml chloramphenicol and isopropyl-*β*-d-thiogalactopyranoside (IPTG) 1 mM or 500 μM when appropriate.

*Escherichia coli* strains were grown at 37°C in LB medium supplemented with 100 μg/ml ampicillin and 20 μg/ml kanamycin. *E. coli* strains DH5α were used to maintain plasmids used for transfection in HeLa cells.The WT amoeba *D. discoideum* strain Ax2 (DBS0235534) was obtained from the Dicty Stock Center (http://dictybase.org). *D. discoideum* cells were grown axenically in HL5 medium at 22°C with 100 μg/ml streptomycin and 66 μg /ml penicillin G. The *A. castellanii* environmental isolate (gift from P. Pernin from Laboratory of Pharmacy, Université Lyon 1, Lyon, France) were grown axenically in PYg medium at 30°C with 100 μg/ml streptomycin and 66 μg /ml penicillin G.

HeLa cells (gift from INSERM U1111, Lyon, France) were maintained at 37°C in 5% CO_2_ in DMEM (Dulbecco’s modified Eagle’s medium) supplemented with 10% heat-inactivated fetal calf serum (FCS). U937 monocyte cells were maintained at 37 °C in 5 % CO_2_ in RPMI 1640 medium (ThermoFisher Scientific) supplemented with 10 % heat-inactivated foetal calf serum (HyClone™). U937 monocytes differentiation into macrophages is conducted during 2 days at a phorbol 12-myristate 13-acetate (PMA) concentration of 100 ng/ml.

### General cloning techniques

The plasmids and primers used in this study are shown in Supplementary Data S2 and S3. Gateway cloning technic was performed for cloning into mammalian expression vectors as recommended by the manufacturer (Invitrogen). SLIC cloning was performed following the procedure described by Li & Elledge^20^ for cloning into bacterial expression vectors. Restriction enzymes, T4 DNA ligase and T4 DNA polymerase were purchased from New England Biolabs. Plasmid DNA from *E. coli* was extracted by Plasmid Midi and Mini Kits (Omega). PCR amplifications were carried out with PrimeStar polymerase as recommended by the manufacturer (Takara). *E. coli* competent cells are transformed by thermal shock with 100 ng of plasmid DNA and *L. pneumophila* strains are transformed by electroporation (2,4 kV, 100 Ω and 25 μF) with 2 μg of plasmid DNA.

DNA fragments corresponding to the *legK2* (*lpp2076*) and *vipA* (*lpp0457*) coding sequences were amplified by PCR using genomic DNA of *L. pneumophila* Paris as a template and specific primers as described in Supplementary Data S3. The coding sequences were inserted into the Gateway pDONR207 vector (Invitrogen) by *in vitro* recombination. The *legK2* and *vipA*-encoding genes were transferred by Gateway cloning from pDONR207-*legK2* and pDONR207-*vipA* to vectors pDEST27 and peGFP (Invitrogen) to produce GST-tagged LegK2 and C or N-terminal GFP-fused VipA proteins in mammalian cells, respectively. The coding sequences amplified by primers were also inserted into the XmaI-linearized pXDC61 vector by SLIC cloning to produce β-Lactamase-fused LegK2 or VipA proteins.

### Gene inactivation and mutation in *L. pneumophila*

A homologous recombination strategy was performed as previously described^21^ in order to obtain *L. pneumophila* Paris mutant strains for *lpp2076* and *lpp0457* genes. The 2000-bp upstream and downstream regions of the gene of interest were amplified by PCR with primers carrying 30-nt sequences (P1-P2 primers pair for the upstream region; P3-P4 primers pair for the downstream region) complementary to a counter-selectable mazF-kan (MK) cassette (Supplementary Data S3). The upstream and downstream regions were assembled to the MK cassette (amplified from plasmid pGEM-mazF-kan with MazFk7-F/MazF-R primers) by PCR overlap extension and used for natural transformation of *Legionella* strains. Transformants were then selected on CYE + kanamycin and counter-selected for sensitivity on CYE + IPTG. Integration of the cassette at the correct locus was also verified by PCR. To obtain scar-free mutants, a second step was performed as follows. Upstream and downstream regions of each gene of interest were amplified with primers carrying a 20-nt tail sequences corresponding to the 3’ end of upstream and downstream region (P1-P5 primers pair for upstream and P4-P6 primer pair for downstream region, Supplementary Data S3), respectively. Theses PCRs were assembled by PCR overlap extension and used to transform the previous Δ*lpp2076*::mazF-kan or Δ*lpp0457*::mazF-kan strains. Transformants were then selected on CYE + IPTG and counter-selected for sensitivity on CYE + Kan. Scar-free deletion of *lpp2076* and *lpp0457* genes were verified by PCR and sequencing.

### Intracellular growth kinetics in *A. castellanii* or U937 macrophages

*L. pneumophila* cells harbouring a fluorescent mCherry protein-producing plasmid were grown on BCYE agar containing 1 M IPTG and 0.5 μg/mL chloramphenicol for 24 hours at 37°C. *A. castellanii* cells were plated in 96-well microplates (10^5^ cells/well) in PY special medium (16 mM MgSO_4_, 40 mM CaCl_2_, 3.4 mM sodium citrate dihydrate, 50 μM Fe(NH_4_)2(SO_4_)2, 2.5 mM Na_2_HPO_4_, 2.5 mM KH_2_PO_4_) and infected at MOI = 5. Infection was synchronized by spinning the infected plates at 2500 rpm for 10 min. Intracellular growth was automatically monitored by measuring mCherry fluorescence at λ_ex_ = 587 nm and λ_em_ = 610 nm on an Infinite M200 microplate reader (Tecan) or by lysing infected amoebae with 0.8 % saponin, plating lysates at different dilutions on BCYE agar plates and counting grown bacteria at day 4 post-plating.

U937 macrophages were plated and differentiated in 96-well microplates (10^5^ cells/well) in RPMI + 10% FBS + PMA medium and infected at MOI = 1. Infection was synchronized by spinning the infected plates at 2500 rpm for 10 min. Intracellular growth was monitored by lysing infected macrophages with sterile water, plating lysates at different dilutions on BCYE agar plates and counting grown bacteria at day 4 post-plating.

### *D. discoideum* infection by *L. pneumophila* for microscopic analysis

*D. discoideum* cells were plated at a concentration of 5×10^5^ cells/mL on sterile glass coverslips a day before infection in MB medium (7.15 g/L yeast extract; 14.3 g/L peptone; 20 mM MES and buffered at a pH 6.9) and incubated overnight at 22°C. Monolayers were infected the next day at an MOI of 100 with mCherry-expressing bacteria grown overnight at 37°C in AYE 1X medium supplemented with chloramphenicol and IPTG for maintenance and induction of mCherry expression plasmid pXDC50. Plates were then spun at 2000 rpm for 10 min and incubated at 25°C for a specific time of infection and further treated for microscopic analysis.

### v-ATPase visualization on LCVs in *D. discoideum* during infection by *L. pneumophila*

*D. discoideum* cells plated on sterile glass coverslips were infected as described above and incubated at 25°C for 1 hour. Monolayers were fixed with 4% formaldehyde, permeabilized with 0.1 % Triton X-100 for 5 min at RT and blocked in 0.2% BSA for 1h at RT. v-ATPase was then stained with anti-VatA antibodies (gift of F. Letourneur, Montpellier, France) and detected with anti-mouse secondary antibody from goat conjugated with the fluorochrome Alexa Fluor 488 (A11029; Molecular Probes). Glass slides were then mounted on slides with Fluoromount (Thermofisher) and microscopy was carried out using a confocal laser scanning microscope (LSM800; Zeiss).

### Actin polymerization in *D. discoideum* during infection by *L. pneumophila*

*D. discoideum* cells plated on sterile glass coverslips were infected as described above and incubated at 25°C for 15 min. Monolayers were fixed with 4% formaldehyde, permeabilized with 0.1 % Triton X-100 for 5 min at RT and blocked in 0.2% BSA for 1h at RT. Actin was then stained with phalloidin-FITC (P5282; Sigma). Glass slides were then mounted on slides with Fluoromount (Thermofisher) and microscopy was carried out using a confocal laser scanning microscope (LSM800; Zeiss).

### RNA isolation, depletion of rRNA and RNAseq

*L. pneumophila* Paris WT strain was grown in AYE medium and harvested by centrifugation (5 min, 7000 rpm, 4°C) at different growth phases: at the exponential phase (optical density of 1.5 at 600nm (OD_600nm_ 1.5)), post-exponential phase (OD_600nm_ 4 and visual check of motility acquisition) and to the onset of stationary growth phase (OD_600nm_ 5). Total RNA from bacterial cultures was extracted accorded to a previously described procedure^22^. Briefly, pellets of 10^9^ bacterial cells were lysed in 50 μl of RNAsnap buffer (18 mM EDTA, 0.025% SDS, 95% formamide) and total RNAs were extracted using a tri-reagent solution (acid guanidinium thiocyanate–phenol–chloroform) and isopropanol-precipitated. After precipitation, we performed an additional step of RNA purification on silica-based columns (DirectZol kit, ZymoResearch) by following the manufacturer’s recommendations. RNA sample purity and concentration were determined by spectrophotometric analysis on a NanoDrop 2000 UV-Vis Spectrophotometer (Thermo). RNA sequencing was performed following ribosomal RNA depletion and cDNA libraries preparation on an NovaSeq platform (Illumina) with paired-end 150bp (Genewiz-Azenta, Leipzig, Germany). After mapping sequence reads to the reference genome and extraction of gene hit counts, the comparison of gene expression between the defined groups of samples was performed using DESeq2.The BAM files were imported into IGV software (V2.15.2) and reads were aligned with the genome sequence of *L. pneumophila* Paris strain (NCBI accession number: NC_006368). We used IGV to visualize data as a graphical display to compare the transcriptomic data between *legK2* and *vipA* obtained from the different experiments.

### Sample preparation and mass spectrometry

Three independent cultures of *L. pneumophila* Paris WT were made in liquid medium AYE at 30°C until reaching OD_600nm_ 1, 2, 3, 4 and 5. Then, a pellet of 3.10^9^ bacteria were made for each independent culture and stored at - 80°C for 24 hours. The cells of each pellet were then lysed by adding 200 μl of B-PER (Bacterial Protein Extraction Reagent) (ThermoScientific) followed by a 15-minutes incubation at 37°C and a 30-minutes centrifugation at 21 000 g at 4°C. The clear lysates were then recovered, and concentration was determined by Bradford assay using the Coomassie Plus kit (Thermo Scientific) according to the manufacturer’s instructions. For each sample, 50 μg of proteins were precipitated with TCA. Pellets were washed, dried and resolubilised in NaOH 50 mM/HEPES 1M pH 8/H_2_O-15/5/78-v/v/v, reduced with 5 mM TCEP for 45 minutes at 57°C, and then alkylated with 10 mM iodoacetamide for 30 minutes in the dark at room temperature and under agitation (850 rpm). Double digestion was performed with endoproteinase Lys-C (Wako) at a ratio 1/100 (enzyme/proteins) in TEAB 100 mM for 5h, followed by an overnight trypsin digestion (Promega) at a ratio 1/100 (enzyme/proteins). Both LysC and Trypsin digestions were performed at 37°C. Peptide concentration was checked before TMT labeling with quantitative fluorometric peptide assay (ThermoScientific). Each sample was labelled with TMTpro 16 plex (ThermoScientific) according to the manufacturer’s instructions. This resulted in 15 samples (3 for each OD), plus one sample “pool” where all samples were mixed in equal quantity before labelling. Then 2 μg of each of the 16 labeled samples were mixed and dried up. The pellet was resuspended in 300 μl 0,1% TFA and fractionated on High pH reserved-phase fractionation spin-column (Thermo Scientific) according to the manufacturer’s instructions for TMT-labelled peptide samples. Recovered fractions were dried up and then resuspended in 10 μl 2% ACN + 0,1% formic acid. LC-MSMS analysis. All fractions were then analysed in triplicate by mass spectrometry (Q Exactive HF coupled with nanoRSLC Ultimate 3000, ThermoScientific). 1 μL of each fraction were injected and loaded on a C18 Acclaim PepMap100 trap-column 300 μm ID x 5 mm, 5 μm, 100Å, (ThermoScientific) for 3 min at 20 μL/min with 2% ACN, 0.05% TFA in H_2_O and then separated on a C18 Acclaim Pepmap100 nano-column, 50 cm x 75 mm i.d, 2 mm, 100 Å (ThermoScientific) with a 60 minutes linear gradient from 3.2% buffer A to 40% buffer B (A: 0.1% FA in H2O, B: 0.1% FA in ACN) and then from 40 to 90% of B in 2 min, hold for 10 min and returned to the initial conditions in 1 min for 14 min. The total duration was set to 90 minutes with a flow rate of 300 nL/min and the oven temperature was kept constant at 40°C. Labelled peptides were analysed with TOP15 HCD method: MS data were acquired in a data dependent strategy selecting the fragmentation events based on the 15 most abundant precursor ions in the survey scan (375-1800 Th). The resolution of the survey scan was 120,000 at m/z 200 Th and for MS/MS scan the resolution was set to 45000 at m/z 200 Th. The Ion Target Value for the survey scans in the Orbitrap and the MS/MS scan were set to 3E6 and 1E5 respectively and the maximum injection time was set to 50 ms for MS scan and 100 ms for MS/MS scan. Parameters for acquiring HCD MS/MS spectra were as follows; collision energy = 32 and an isolation width of 1.2 m/z. The precursors with unknown charge state, charge state of 1 and 8 or greater than 8 were excluded. Peptides selected for MS/MS acquisition were then placed on an exclusion list for 30 s using the dynamic exclusion mode to limit duplicate spectra. Data were analysed using Proteome Discoverer 2.4 with the SEQUEST HT search engine on the *L. pneumophila* genome from NCBI (NC_006368) and a database of common contaminants. Precursor mass tolerance was set at 10 ppm and fragment mass tolerance was set at 0.02 Da, and up to 2 missed cleavages were allowed. Oxidation (M), acetylation (Protein N-terminus) and phosphorylation (S, T, Y) were set as variable modification and TMTpro labelled peptides in primary amino groups (K and N-ter), and Carbamidomethylation (C) as fixed modification. Validation of identified peptides and proteins was done using a target decoy approach with a false positive (FDR < 1%) via Percolator. Protein quantitation was performed with reporter ions quantifier node in Proteome Discoverer 2.4 software with integration tolerance of 20 ppm, peptide and protein quantitation based on pairwise ratios and t-test hypothesis test. Protein expression of RocC, VipA and LegK2 were extracted from the obtained dataset (see Supplementary Data S7). RocC was used as a control as its expression has been extensively studied via classical methods (Western-Blot)^23^. LegK2 was not detected.

### TEM Translocation assays

U937 cells grown in RPMI supplemented with 10% FCS were plated in black clear-bottom 96-well plate at a 1×10^5^ cells/well concentration in presence of 100 ng/mL of Phorbol-12-myristate-13-acetate (PMA) to allow differentiation of U937 cells in macrophages. Overnight cultures of *L. pneumophila* strains carrying either pXDC61-*legK2* or pXDC61-*vipA* were grown in AYE + 5 μg/mL chloramphenicol and 500 μM IPTG to induce the production of TEM-fused proteins. Bacterial suspension in RPMI at 2×10^8^ cells/mL were used to infect U937 cells (MOI = 50). After centrifugation at 2500 rpm for 10 min to initiate bacteria-cell contact, the infected cells were incubated at 37°C with 5% CO_2_. At different time points, 10 μM of CCCP were added to block effector translocation through the Dot/Icm T4SS. Cell monolayers were then loaded with the fluorescent substrate by adding 20 μl of 6X CCF4/AM solution (LiveBLAzer-FRET B/G Loading Kit, Invitrogen) containing 15 mM Probenecid (Sigma). The cells were incubated for an additional 90 min at room temperature. Fluorescence was quantified on an Infinite M200 microplate reader (Tecan) with excitation at 405 nm (10 nm band-pass), and emission was detected via 460 nm (40 nm band-pass, blue fluorescence) and 530 nm (30 nm band-pass, green fluorescence) filters.

### Protein localization in transfected mammalian cells

HeLa cells were plated one day before transfection on sterile glass coverslips and transfected or cotransfected with empty pDEST27 or pDEST27-*legK2*, empty peGFP or peGFP-N-VipA and/or pCI-Neo3Flag-ARPC1B using JetPrime (Polyplus). At 24h post-transfection, cells were fixed with 4% formaldehyde, quenched with 0,1 μg/ml glycin, permeabilized with 0.3% Triton X-100 and blocked with 1% BSA. GST and GST-LegK2 proteins were labelled with an anti-GST antibody from rabbit (A7340; Sigma) and detected with an anti-rabbit secondary antibody from goat conjugated with the fluorochrome Alexa Fluor 488 (A11034; Molecular Probes) or fluorochrome Alexa Fluor 594 (A11037; Molecular Probes). Flag-ARPC1B were labelled with anti-Flag antibody from mouse (F1804; Sigma) and detected with anti-mouse secondary antibody from goat conjugated with the fluorochrome Alexa Fluor 594 (A11032; Molecular Probes). EEA1 endogenous protein was labelled with an anti-EEA1 antibody from rabbit (3288S; CellSignalling Technology). Glass slides were then mounted on slides with Fluoromount and microscopy was carried out using a confocal laser scanning microscope (LSM800; Zeiss).

### Protein localization in *D. discoideum* during infection by *L. pneumophila*

*D. discoideum* cells were plated on sterile glass coverslips a day before infection in MB medium (7.15 g/L yeast extract; 14.3 g/L peptone; 20 mM MES and buffered at a pH 6.9) and incubated overnight at 22°C. Monolayers were infected the next day at an MOI of 50 with HA-protein -expressing bacteria grown overnight at 37°C in AYE 1X medium supplemented with chloramphenicol and IPTG for maintenance and induction of HA expression plasmid. Plates were then spun at 2000 rpm for 10 min and incubated at 25°C. At different time points, Monolayers were fixed with 4% formaldehyde, permeabilized with 0.1 % Triton X-100 for 5 min at RT and blocked in 0.2% BSA for 1h at RT. HA-fusion protein were stained with anti-HA primary antibodies and then with anti-mouse-coupled to Alexa Fluor 488 antibodies (A-11029; ThermoFisher Scientific). Glass slides were then mounted on slides with Fluoromount + DAPI (Thermoscientific) and microscopy was carried out using a confocal laser scanning microscope (LSM800; Zeiss).

### Co-immunoprecipitation by GFP-Trap

HeLa cells were plated in 10-cm Petri Dish the day before transfection in DMEM supplemented with 10% fetal calf serum FCS. Next day, they were co-transfected with either pDEST27 or pDEST27-*legK2*, empty peGFP, peGFP-C-VipA or peGFP-N-VipA using JetPirme (Polyplus). At 24h post-transfection, cells were harvested in ice-cold PBS, washed and pelleted cells were lysed during 1h in ice-cold RIPA buffer (10 mM Tris, pH 7.5, 150 mM NaCl, 0.5 mM EDTA, 0.1% SDS,1% Triton X100, 1% deoxycholate, benzonaze, 2,5 mM MgCl_2_, 1 mM PMSF and protease inhibitors) at 4°C with gentle agitation. Lysed extracts were then centrifuged at 20 000 g for 10 min at 4°C. Lysates were diluted in washing buffer and then incubated with washed GFP-Trap® Agarose beads (Chromotek) for 1h30. After incubation, beads were collected by centrifugation, washed three times before eluting GFP-tagged proteins and their potential interactants in 100 μL of Laemmli buffer 2X at 95°C for 7 min.

### Overexpression and purification of GST-LegK2 and 6His-VipA

*E. coli* BL21 (DE3) strains carrying either pGEX-*legK2* or pQE30-*vipA* were grown at 37°C until cultures reached an OD_600 nm_ of 0.7. Then, IPTG was added to a final concentration of 0,2 mM and growth was continued overnight at 20°C. Pellets were resuspended in GST-Pull Down equilibration/wash buffer (125 mM Tris-HCl pH 7.5, 150 mM NaCl + Protease Inhibitor Cocktail (Sigma) + 1 mg/ml lysozyme) or in 6His-Pull Down equilibration/wash buffer (50 mM Tris-HCl pH 7.5, 150 mM NaCl and 10 mM Imidazole + Protease Inhibitor Cocktail (Sigma) + 1 mg/ml lysozyme) and bacteria were lysed using 3 passages in a French Press (SLM, Urbana, IL). After lysate centrifugation at 12 000g for 30 min, supernatants were collected and transferred to tubes containing either Pierce™ Gluthatione Magnetic Agarose Beads (ThermoFisher Scientific) or TALON Metal Affinity Resin (Takara Bio) respectively, according to the manufacturers’ recommendations. The purity of the eluted protein was analysed by SDS-PAGE.

### *In vitro* phosphorylation assays

*In vitro* phosphorylation of 2 μg of purified 6His-VipA fusion protein was performed for 30 min at 37°C in 20 μl of a buffer containing 25 mM Tris-HCl pH 7.5, 5 mM MnCl2, 5 mM dithiothreitol, 100 mM ATP. 1 μg of myelin basic protein (MBP) was added as positive phosphorylation control for LegK2. In each case, the reaction was stopped by the addition of an equal volume of 2X Laemmli loading buffer. Proteins were then separated by SDS-PAGE and immunoblotted with an anti-phosphothreonine monoclonal antibody (CST; #9381).

### Phylogenetic profiles of VipA and LegK2

A total of 647 annotated *Legionella sp*. and *Coxiella burnetti* genomes were downloaded from NCBI as of October 2017 (accession numbers are available in Supplementary Data S4). We used MMSEQS2 (default parameters)^24^ to cluster the annotated proteins into families. 95 universal-unicopy family were aligned with MAFFT (default parameters)^25^ and concatenated at the nucleotide level to reconstruct a phylogenetic tree using FastTree2.1^26^. We used Treemmer^27^ to select 120 genomes representing the phylogenetic diversity in this tree, with the constraint that the 19 *L. pneumophila* genomes missing either VipA or LegK2 were represented. This tree was then used in Count^28^, with the complete table of presence/absence of proteins in the corresponding genomes to infer the parameters of the probabilistic model of Gain-Duplication-Loss. These parameters were then used to reconstruct the ancestral presence/absence of all genes. We finally used iTOL^29^ to represent the evolutionary history of VipA and LegK2.

## RESULTS

### LegK2/VipA effector pair controls actin polymerization at the surface of the LCV

Taking into account the antagonistic activities of LegK2 and VipA towards actin polymerization, and the localization and role of LegK2 in the inhibition of actin polymerization on the LCV, we hypothesized that LegK2/VipA may cooperatively contribute to the remodeling of actin cytoskeleton at the surface of the LCV during *Legionella* infection.

Scar-free single mutants Δ*legK2*, Δ*vipA*, and double mutant Δ*legK2/*Δ*vipA* of *L. pneumophila* Paris strain were constructed in two steps, taking advantage of a homologous-recombination strategy with the counter-selectable *mazF-kan* cassette, as previously described^21^. *Dictyostelium discoideum* was infected with mCherry-producing *L. pneumophila* Paris strain, Δ*dotA*, Δ*legK2*, Δ*vipA* single mutants, or Δ*legK2/*Δ*vipA* double-mutant strains, and polymerized actin was visualized with phalloidin-FITC, 15 min post-infection **(figure 1A)**. While less than 5% of WT bacteria containing vacuoles were labeled with phalloidin, more than 20% of the avirulent *dotA* mutant containing vacuoles were actin-positive **(figure 1B)**. This significative difference highlights the importance of local actin remodeling on the LCV during *Legionella* infectious cycle. Indeed, less than 1 min after engulfment of the bacteria, cortical actin associated with the bacterial entry sites during phagocytosis dissociates from the LCV^30^. Similarly to what was reported for the Lens strain^16^, 18% of Δ*legK2* Paris mutant LCVs were labeled with phalloidin, thus confirming that LegK2 plays a key role in the inhibition of actin polymerization at the surface of the LCV **(figure 1B)**. In contrast, the contribution of VipA in controlling actin polymerization on the LCV surface is limited or non-detectable, as only 3.5% of LCVs are decorated with polymerized actin in the Δ*vipA* mutant, which is not significantly different from the parental Paris strain. Nevertheless, this activity is able to compensate the actin polymerization inhibition defect caused by the absence of LegK2, as 5% of the vacuoles containing the Δ*legK2/*Δ*vipA* double mutant are actin-positive, which is not significantly different from the parental strain but significantly different to that observed in the Δ*legK2* mutant **(figure 1B)**.

**Figure 1.**
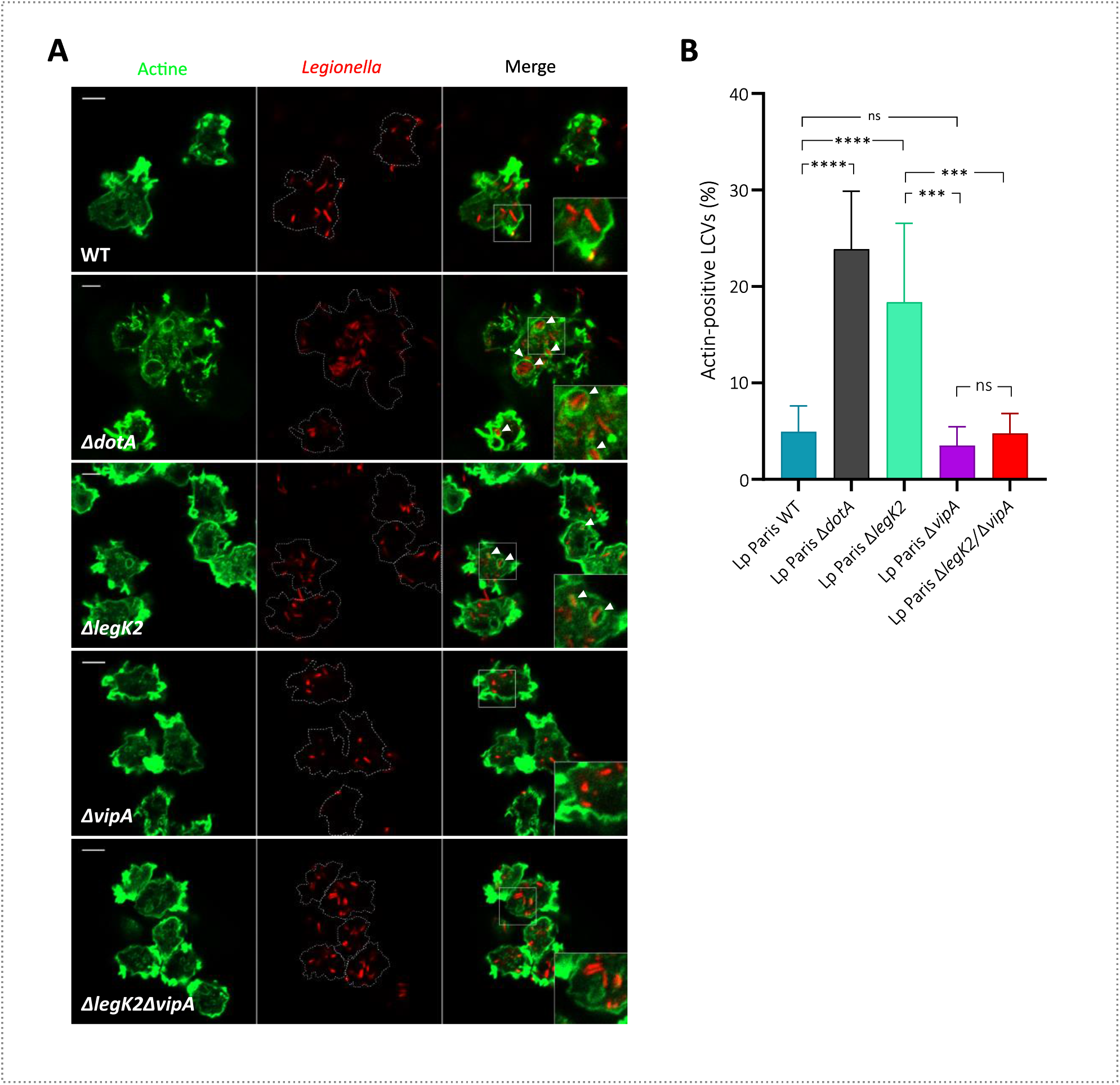
LegK2/ VipA effector pair controls actin polymerization at the surface of the LCV. **(A)** Actin polymerization on LCVs during *Legionella* infection. *D. discoideum* was infected for 15 min at an MOI of 100 with mCherry-labelled WT, Δ*dotA-*, Δ*legK2*-, Δ*vipA-* and Δ*legK2*/Δ*vipA*- mutant *L. pneumophila* strains. Polymerized actin on LCVs was detected by labelling with phalloidin-FITC. White arrows show examples of actin-positive LCVs. Scale bar, 10 μm. **(B)** Detection of polymerized actin in Δ*dotA*, Δ*legK2*, Δ*vipA*, Δ*legK2/ΔvipA* mutant-containing vacuoles. Actin-positive vacuoles (*n* >100) were counted for amoeba infected with *L. pneumophila* WT, the derivative Δ*dotA*, Δ*legK2*, Δ*vipA*, Δ*legK2/ΔvipA* mutants. Quantification data are representative of three independent experiments, and the error bars represent the standard deviations from triplicates. Statistical analyses were performed using two-way ANOVA test for intracellular replication and one-way ANOVA test for v-ATPase detection on LCVs: ns, so significant difference; *, *p* < 0.05; **, *p* < 0.005; ***, *p* < 0.0005; ****, *p* < 0.0001.

Taken together, these data point out that deletion of *vipA* restores the Δ*legK2* mutant defect in controlling actin polymerization at the surface of the LCV, thus demonstrating that the LegK2/VipA effector pair cooperatively controls actin polymerization during *L. pneumophila* infection.

### Deletion of *vipA* suppresses the endosomal escape and intracellular replication defects of the Δ*legK2* mutant

In addition to sharing antagonistic activities towards actin polymerization, both LegK2 and VipA target the host cell endosomal pathway. Specifically, LegK2 has been shown to inhibit actin polymerization at the LCV surface and subsequent recruitment of late endosomes/lysosomes to the vacuole surface^16^, and VipA is an actin nucleator that colocalizes with early endosomes and has been proposed to disrupt normal vacuolar trafficking pathways in host cells^14^. In this context, we addressed the question of the functional relationship between these two effectors regarding phagosome maturation, in particular their contribution to the inhibition of LCV fusion with late endosomes. *D. discoideum* was infected with mCherry labeled *L. pneumophila* Paris strain WT, Δ*dotA*, single Δ*legK2-* or Δ*vipA* mutants or the double Δ*legK2/*Δ*vipA* mutant, and the late endosomal or lysosomal vacuolar H^+^-ATPase (v-ATPase) was labelled 1 h post-infection by immunofluorescence with anti-VatA antibodies **(figure 2A)**. As expected, the Δ*dotA* mutant is unable to block phagosome maturation and shows much more VatA-positive vacuoles (38%) compared to a WT strain (12,8%) **(figure 2B)**. Reminiscent to the Δ*dotA* strain, the Δ*legK2* mutant presents about 36% of VatA-positive vacuoles, which confirms that LegK2 plays a key role in the inhibition of phagosome maturation **(figure 2B)**. Interestingly, while only 13% of vacuoles were VatA-positive for the Δ*vipA* strain (which indicates that VipA is not necessary to inhibit phagosome maturation), the Δ*legK2/*Δ*vipA* double-mutant also shows 12% of VatA-positive vacuoles which is not significantly different from the WT-containing vacuoles **(figure 2B)**. Thus, the Δ*vipA* mutation is able to compensate for the phagosome maturation defect caused by the Δ*legK2* mutation.

**Figure 2.**
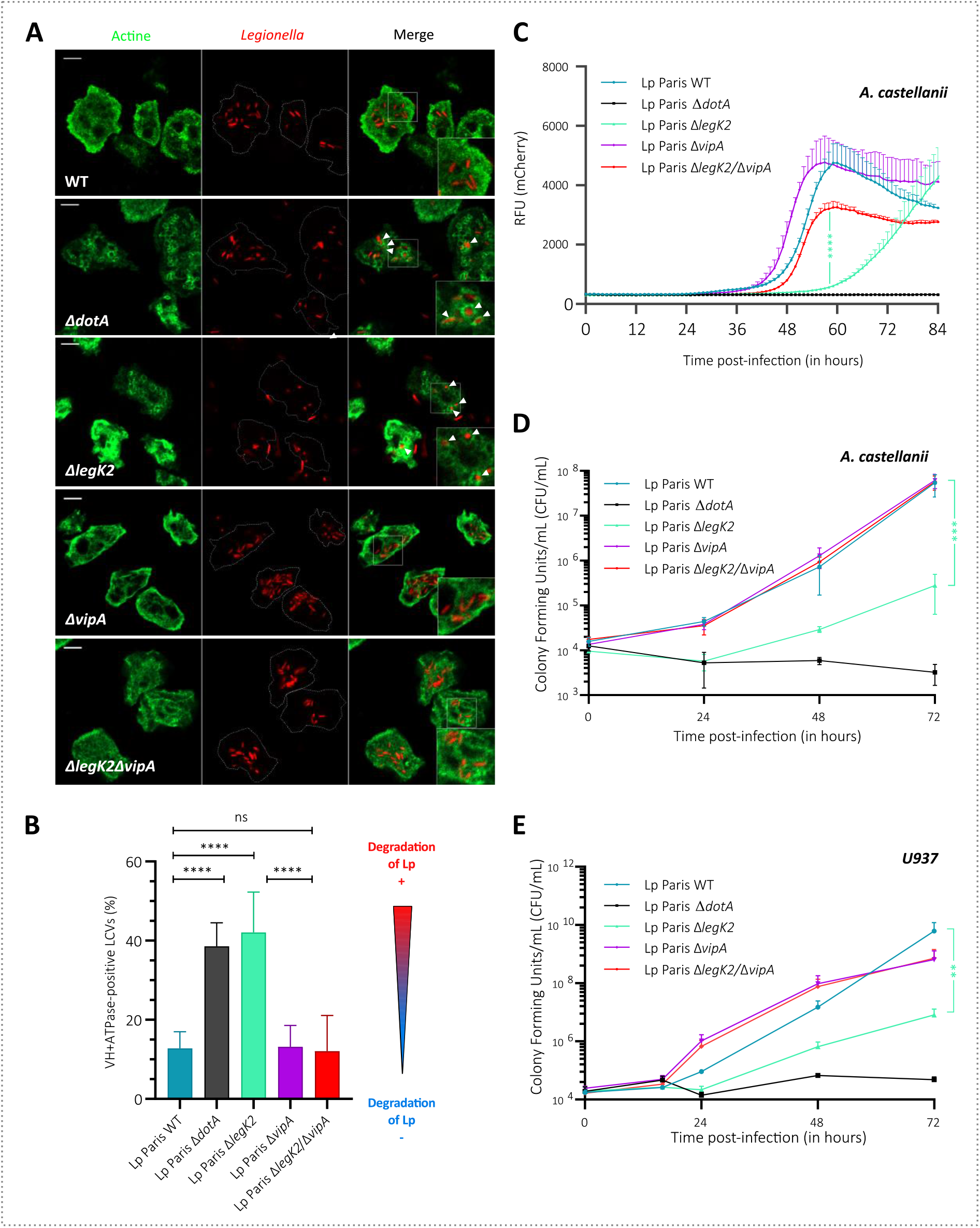
Deletion of *vipA* suppresses the endosomal escape and intracellular replication defects of the Δ*legK2* mutant. **(A)** Acquisition of vacuolar proton ATPase on LCVs. *D. discoideum* was infected for 1 h at an MOI of 100 with mCherry-labelled WT, Δ*dotA-*, Δ*legK2*, Δ*vipA-* and Δ*legK2*/Δ*vipA*-mutant *L. pneumophila* strains. The presence of vacuolar H+-ATPase on LCVs was detected by an immunofluorescence assay with anti-VatA antibodies. **(B)** Detection of vacuolar proton ATPase on LCVs containing *L. pneumophila* Paris WT strain, Δ*dotA*, Δ*legK2*, Δ*vipA*, Δ*legK2/ΔvipA L. pneumophila* mutants. VatA-positive vacuoles (*n>1*00) in amoebae infected with *L. pneumophila* WT, the derivative Δ*dotA*, Δ*legK2*, Δ*vipA*, Δ*legK2/ΔvipA*, mutants were counted. These data are representative of three independent experiments, and the error bars represent the standard deviations from triplicates. Statistical analyses were performed using two-way ANOVA test for intracellular replication and one-way ANOVA test for v-ATPase detection on LCVs: ns, no significant difference; *, *p* < 0.05; **, *p* < 0.005; ***, *p* < 0.0005; ****, *p* < 0.0001. **(C)** Intracellular replication of *L. pneumophila* in *A. castellanii* measured by fluorescence. *A. castellanii* amoebae were infected at MOI = 5 with *L. pneumophila* Paris WT strain or *ΔdotA, ΔlegK2, ΔvipA, ΔlegK2/ΔvipA* mutant *L. pneumophila* strains transformed with mCherry-expressing plasmids and intracellular replication was monitored by quantifying mCherry-fluorescence intensity. **(D and E)** Intracellular replication of *L. pneumophila* in *A. castellanii* measured by CFU (D) and in U937-derived macrophages (E). *A. castellanii* amoebae and U937-derived macrophages were infected at MOI = 5 and MOI = 1 respectively, with *L. pneumophila* Paris WT strain, Δ*dotA*, Δ*legK2*, Δ*vipA*, Δ*legK2/ΔvipA* mutant *L. pneumophila* strains and intracellular replication was monitored by numbering CFU (colony forming units) after amoeba/macrophages lysis at different times post-infection on BCYE agar medium. Results shown in C, D and E are representative of three independent experiments, and the error bars represent the standard deviations from triplicates. Statistical analyses were performed using two-way ANOVA test for intracellular replication and two-way ANOVA test: ns, so significant difference; *, *p* < 0.05; **, *p* < 0.005; ***, *p* < 0.0005; ****, *p* < 0.0001

The ability of *L. pneumophila* to escape endosomal degradation is the prerequisite for its intracellular replication. Therefore, we sought to identify whether the LegK2/VipA functional interaction could impact *L. pneumophila* intracellular replication. *L. pneumophila* Paris strain or its mutant derivatives expressing the mCherry fluorescent protein on a plasmid were used to infect the amoeba *Acanthamoeba castellanii* at a MOI of 5. Bacterial intracellular growth was monitored by fluorescence measurement during 84h **(figure 2C)**. As expected, wild-type *L. pneumophila* started efficient intracellular growth at 40 h post-infection, while the T4SS Δ*dotA* mutant failed to replicate. The Δ*vipA* mutant showed intracellular multiplication from the same time with the same growth rate compared to the Paris strain, while the Δ*legK2* mutant was significantly delayed for intracellular multiplication, as previously reported for Lens strain^31^, thus confirming the key role of this effector in the virulence of *L. pneumophila*. More interestingly, the deletion of *vipA* fully complements the intracellular multiplication defect of the Δ*legK2* mutant, revealing the first example of effector-effector functional interaction with strong impact on *L. pneumophila* virulence **(figure 2C)**. The functional complementation of Δ*legK2* mutant by the Δ*vipA* deletion was confirmed by numbering the CFU resulting from the lysis of *A. castellanii* infected by Δ*legK2*, Δ*vipA*, or double mutant Δ*legK2*/Δ*vipA* strains **(figure 2D)**. The same phenotype was observed upon infection of U937 macrophages **(figure 2E)**. Importantly, full genome sequencing of Δ*legK2*, Δ*vipA*, and Δl*egK2/*Δ*vipA* strains revealed no other secondary mutations, confirming that the complementation of the intracellular replication defect of Δ*legK2* strain is solely due to the deletion of the *vipA* gene **(Supplementary Data S5)**. It is noteworthy that none of the single and double mutants show an axenic growth defect **(Supplementary Data S6)**.

Together, these data identify the first effector-effector suppression pair targeting the host cell actin cytoskeleton and show that LegK2/VipA pair contributes to bacterial escape from the endosomal pathway and its subsequent intracellular replication. They also demonstrate that VipA contributes to the control of vacuolar endosomal trafficking, consistent with its localization with early endosomes in transfected cells, thus revealing its role in the *L. pneumophila* infectious cycle, despite the absence of a defect of the single *vipA* deleted mutant.

### LegK2 and VipA effectors are produced and secreted at the early stage of infection

To investigate in detail the molecular relationship between LegK2 and VipA effectors, we addressed their expression/secretion pattern. First, we performed an RNA-seq from *L. pneumophila* Paris strain grown in nutrient-rich medium to exponential (Expo; OD_600_ 1.5), post-exponential (Post-Expo; OD_600_ 4 and visual check of motility acquisition) and to the onset of stationary (Stat; OD_600_ ∼6) growth phase. Read counts mapped on *L. pneumophila* Paris genome show that both *vipA* and *legK2* genes are weakly expressed. Nevertheless, while *vipA* mRNA is overexpressed in the exponential growth phase compared to the other two phases **(figure 3B)**, *legK2* mRNA reads are mostly detected in post-exponential and stationary phases **(figure 3A)**. These results are in accordance with other RNAseq analysis available in the literature and realized by Sahr et al.(2012)^32^. In addition, the relative amount of *L. pneumophila* Paris WT proteome at different OD during growth in nutrient-rich medium was analyzed, by a high-resolution quantitative mass spectrometry approach **(figure 3B)**. Of note, the RocC protein used as a control was detected by this approach according to a pattern similar to the one previously obtained by Western-blot^23^. VipA was detected accumulating from OD_600_ = 1 to OD_600_ = 4 before diminishing at OD_600_ = **5 (figure 3B; Supplementary Data S7)**. As the RNAseq data showed that the gene was more expressed during the exponential phase, these mass spectrometry data suggest an unreported post-transcriptional control that results in the overproduction of VipA during the post exponential growth phase. Consistent with the low level of l*egK2* mRNA, the LegK2 protein was not detected at any growth phase, confirming its low expression level. Interestingly, these latter results are in agreement with multi-genome microarray experiment performed during *A. castellanii* infection with *L. pneumophila* Paris strain, which revealed that *legK2* is 1.94-fold upregulated at 14h post-infection compared to 8h post-infection and that *vipA* is 1.21-fold upregulated 14h post-infection compared to 8h post-infection^33^. Together, these data suggest that the LegK2 and VipA effectors are produced in the transmissive phase of the *Legionella* infectious cycle, and therefore are available for secretion right after the contact with the host cell, at the early stage of the infection cycle.

**Figure 3.**
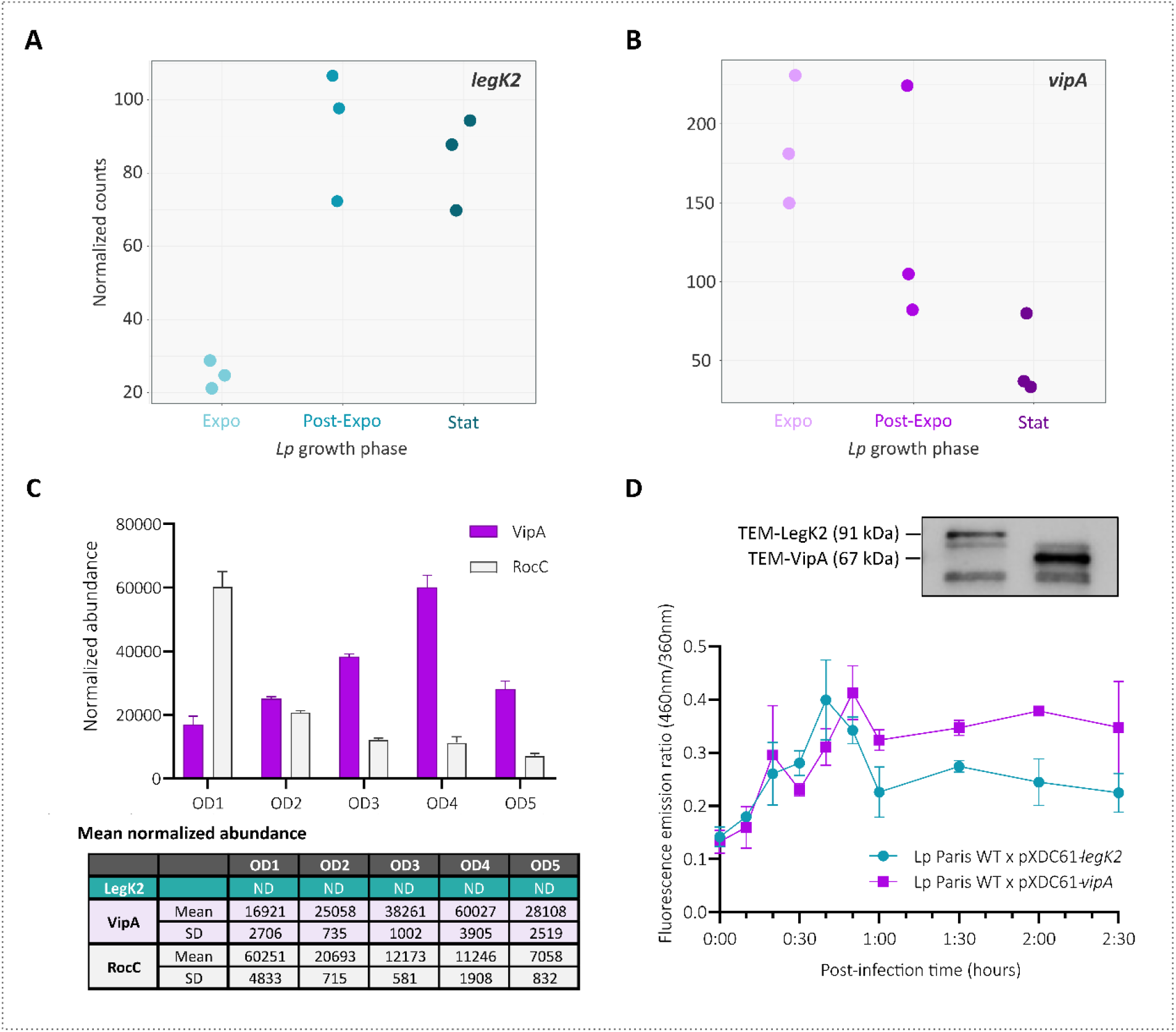
LegK2 and VipA effectors are produced and secreted at the early stage of infection. **(A and B)** *legK2* and *vipA* genes are weakly expressed, mostly in post-exponential and stationary phases for *legK2* **(A)**, and in the exponential growth phase for *vipA* **(B)**. Three cultures of *L. pneumophila* Paris WT were grown in liquid medium AYE (37°C). At the desired growth phases (exponential/OD_600_ = 1.5, post-exponential/start of mobility acquisition/OD_600_ = 4 and stationary collected 2 hours after the post-exponential sample), samples were collected, and their RNA content was analyzed by RNAseq. The graphs show normalized read counts of *legK2* or *vipA* mRNAs for each sample at the different growth phases. **(C)** VipA accumulates in *L. pneumophila* up to DO_600_= 4. Three independent cultures of *L. pneumophila* Paris WT were grown in liquid medium AYE (at 30°C). At the desired OD_600_ (1, 2, 3, 4 and 5), samples were collected, and their protein content was analysed by mass spectrometry after sample-specific labelling (see Material and methods for details). The graphs show the normalized abundance of protein detected at each OD_600_. The LegK2 protein has not been detected in our conditions. The RocC protein is used as a control as it is detected at the same range of quantity as VipA and its production during growth was previously studied by Western-blot ^23^. Of note the RocC pattern of production detected by mass spectrometry corresponds to the one previously obtained by Western blot^23^. **(D)** Translocation kinetics of TEM-LegK2 and TEM-VipA fusion proteins. U937 cells were infected (MOI = 20) with wild-type (WT) Paris strains expressing fusion proteins TEM-LegK2 or a TEM-VipA expression plasmid. Induction of fusion protein expression by 0.5 mM IPTG has been monitored by Western-Blot analysis. Results are obtained from 3 independent experiments made in triplicates and are presented as means ± SD.

Thus, we sought to establish the Dot/Icm translocation kinetics of each effector (independent of its production) by performing VipA and LegK2 kinetic translocation assays using the β-lactamase translocation reporter system^34,35^. U937-derived phagocytes were infected with *L. pneumophila* strains expressing a fusion protein between the TEM-1 β-lactamase and the effector of interest (TEM-LegK2 or TEM-VipA) from a plasmidic IPTG-inducible (Ptac) promoter, as confirmed by western-blot **(figure 3D)**. At different times post-infection, secretion was inhibited by the protonophore CCCP, and levels of secreted effector were quantified by adding CCF4 to the infected cells^36^. CCF4 is composed of coumarin (λ_ex_ =409 nm), and fluorescein linked by a β-lactam ring and fluoresces in green (520nm). The secretion of the fusion protein TEM-effector cleaves the β-lactam ring and induces emission wavelength to change from green to blue (447nm); measuring the blue/green ratio thus reveals the level of secreted effector. Translocated LegK2 effector increases steadily up to 45 minutes post-infection and then decreases and stabilizes **(figure 3D)**. Reminiscent of the secretion profile obtained for LegK2, VipA levels also showed an increase up to 50 minutes post-infection (with a small intermediate peak at 20 minutes post infection) and then a decrease and a stabilization **(figure 3D)**. Together, these data may indicate that LegK2 and VipA effectors are secreted at the same time during the early stage of *L. pneumophila* infection cycle. Noteworthy, the low level of expression of LegK2 does not prejudge the importance of its role during the infectious cycle, since its deletion strongly impairs intracellular replication of *L. pneumophila*.

### The LegK2/VipA suppression pair does not meet the definition of metaeffector

Effector-effector suppression is the hallmark of an emerging class of proteins called metaeffectors, or “effectors of effectors”. The concept of metaeffector, even if remaining flexible, implies a direct physical interaction between an effector and its cognate effector^19^. Taking into account that LegK2 and VipA may be secreted at the same time into the host cell, we studied their localization and possible physical interaction inside eukaryotic cells. Vectors pDEST27-*legK2* and peGFP-N-*vipA* were constructed to express in mammalian cells N-terminal GST-tagged LegK2 (GST-LegK2), and C-terminal GFP-tagged VipA (VipA-GFP), respectively. Sub-cellular localization of GST-LegK2 and VipA-GFP subunits were analyzed after transfection in HeLa cells. ARP2/3 complex and early endosomes were immunolabeled with anti-ArpC1B subunit and anti-EEA1 antibodies, respectively. GST-LegK2 and ArpC1b were detected in the cytoplasm and at the periphery of cells with similar staining patterns, thus suggesting that LegK2 from Paris strain co-localizes with ARPC1B **(figure 4A)**. VipA-GFP and EEA1 appeared as puncta that seem to co-localize as reported in Bugalhão et al. (2016)^37^ **(figure 4A)**. Importantly, co-transfection of Hela cells by p-DEST27-*legk2* and peGFP-N-*vipA* vectors (encoding GST-LegK2 and VipA-GFP) showed the same respective cellular sub-localization to that of LegK2 and VipA when present alone in the cell, demonstrating that each effector does not interfere with the localization of the other one **(figure 4A)**. These data, similar to those previously obtained individually for each effector^14,16,37^, refute the hypothesis of co-localization and localization interference of LegK2 and VipA effectors in transfected mammalian cells.

**Figure 4.**
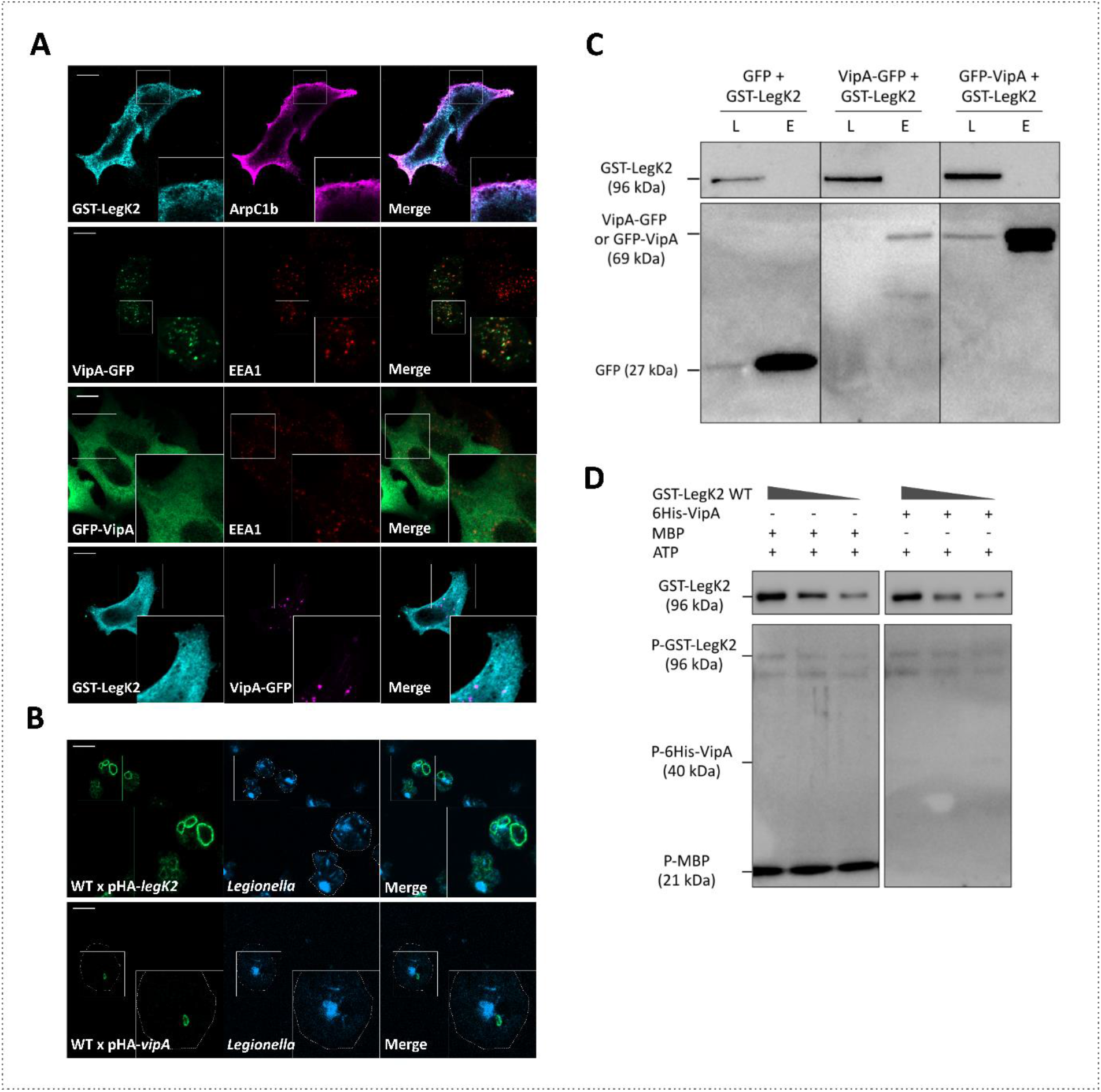
The LegK2/VipA suppression pair does not meet the definition of metaeffector. **(A)** Cellular localization of ARPC1B/LegK2 and EEA1/VipA proteins in HeLa cells transfected by pDEST27-*legK2* or peGFP-N-vipA. GST-LegK2 proteins were detected by immunofluorescence with anti-GST antibodies (Sigma, green), and ARPC1B was detected by anti-ARPC1B antibodies (SantaCruz; red). VipA-GFP proteins were detected thanks to GFP fusion and EEA1 by anti-EEA1 antibodies (CellSignaling Technology). Scale bar represents 10 μm. **(B)** *D. discoideum* were infected at MOI = 50 with *L. pneumophila* WT Paris transformed with pMMB207c-Ptac-HA-*legK2* or pMMB207c-Ptac-HA-*vipA* plasmid. Infected cells were fixed 1-hour post-infection, and HA-tagged proteins were labelled by immunofluorescence with HA-antibodies described (Materials and Methods). DNA was stained with DAPI. Scale = 10 μm. **(C)** GFP-trap co-purification assay of GFP-tagged VipA with GST-tagged LegK2 proteins. HeLa cells were co-transfected by pDEST27 or pDEST27-*legK2* and peGFP-N-VipA or peGFP-C-VipA. GFP or GFP-tagged VipA were purified on GFP-Trap agarose beads and finally, total lysates (L) and eluted fractions (E) were immunoblotted with both anti-GST and anti-GFP antibodies. **(D)** *In vitro* phosphorylation assays of 6his-VipA by LegK2 detected by Western blot assay with antiphosphothreonine antibodies. The 6His-VipA fusion protein purified from *E. coli* BL21(DE3) was incubated with purified GST-LegK2 in the presence of 100 μM ATP. The Myelin Basic Protein (MBP), known to be a substrate of LegK2^31^, was used as a positive control of phosphorylation. Proteins were then separated by SDS-PAGE and detected with antiphosphothreonine antibodies.

Nonetheless, we sought to assess the localization of these effectors during infection. *D. discoideum* were infected at MOI 50 by *L. pneumophila* WT or Δ*dotA* Paris strain transformed by pMMB207c-HA-*legK2* or pMMB207c-HA-*vipA*, and the tagged fused effectors produced upon induction with IPTG were detected in the host cell by anti-tag immunofluorescence at various times post-infection. At one-hour post-infection, HA-LegK2 signal was detected on the LCV surface when cells were infected with the WT Paris strain while no signal was detected upon infection by the Δ*dotA* mutant **(figure 4B; Supplementary Data S7C)**. Conversely, HA-VipA seems to be secreted inside the host cytosol and to form aggregates surrounding the bacteria **(figure 4B)**. The same results were obtained at 30 min, 45min and 1,5 hours post-infection (**Supplementary Data S7A**,**B**), as previously described for experiments conducted up to 8 hours post-infection^14^. Whether the protein is localized at the LCV surface is not clear, especially if comparing with the clear LCV delimitation observed in *D. discoideum* infected with *L. pneumophila* WT expressing HA-LegK2. Yet, VipA-derived peptides were previously identified by mass spectrometry on the LCV surface 1h post-infection from both infected *D. discoideum* and macrophages^38^. The discrepancy between the results of immunodetection and those obtained by mass spectrometry is most likely due to the difference in sensitivity of the two techniques. Noteworthy, the localization of VipA on the surface of the LCV could result from the maturation of the phagocytic vacuole along the endosomal pathway, which involves the fusion of the LCV with early endosomes, with which VipA colocalizes. Altogether, these data suggest that LegK2 and VipA could localize to the LCV surface during infection, at least temporarily, about 1 hour after infection.

Given this temporary common location, we investigated in mammalian cells putative interactions between GST-LegK2 and VipA-GFP by co-affinity purification. HeLa cells were co-transfected with GST-LegK2 and VipA-GFP encoding vectors, and VipA-GFP were purified by GFP-Trap. GFP-tagged VipA protein (67 kDa) was undetectable 24 h post-transfection in the soluble fraction, most likely due to its association within EEA1 membranes, but it was well purified **(figure 4C)**. Although GST-LegK2 (90 kDa) was well expressed and detected in lysates 24 h after transfection, it was not copurified with the VipA fusion protein **(figure 4C)**, suggesting that LegK2 and VipA effectors do not interact in our experimental conditions. To ensure that the low amount of VipA was not the limiting factor for detecting LegK2/VipA interaction, we sought to improve the amount of purified VipA by constructing the peGFP-C-vipA vector encoding the N-terminal GFP-tagged VipA (GFP-VipA). GFP-VipA that localized in the cytosol of transfected cells **(figure 4A)** was well expressed in the soluble fraction and purified efficiently **(figure 4C)**. However, despite the high amount of GFP-VipA, GST-LegK2 was not copurified with the VipA fusion protein **(figure 4C)**. Finally, the phosphorylation status of purified 6His-VipA fusion protein was established by *in vitro* phosphorylation assays and analyzed by Western blot revealed with anti-phosphothreonine antibodies. Phosphorylated form of 6His-VipA protein was not detected in the presence of GST-LegK2 **(figure 4D)**. Consistently, no VipA-derivative phosphopeptides were revealed by our comprehensive proteomic analysis of *L. pneumophila* effector content, at any growth phase, thus suggesting that VipA is not a substrate of the protein kinase LegK2.

Taken together, these data demonstrate that the LegK2/VipA suppression pair does not meet the definition of metaeffector in that these effectors are not capable of interacting with each other, nor is VipA a substrate for phosphorylation of the LegK2 protein kinase. Rather, the relationship between LegK2 and VipA would be an indirect functional antagonism that occurs through counteracting activities on a shared host pathway, namely actin polymerization at the LCV surface, by targeting two distinct cellular partners, ARP2/3 for LegK2 and G-actin for VipA.

### The functional antagonism of LegK2/VipA pair is supported by evolutionary co-occurrence of *legK2/vipA* genes in *L. pneumophila* species

The effector-effector suppression pairs are considered to have evolved to balance the targeting of host cell pathways which, if excessive, could be detrimental to the host and counterproductive for *Legionella* intracellular replication. Besides, effector/metaeffector or other effector-effector suppression pairs are often encoded by adjacent genes on the genome, presumably resulting from the simultaneous acquisition of these genes through horizontal gene transfer. Noteworthy, it is not the case for LegK2 and VipA that are encoded by distant genes on the *L. pneumophila* genome. Thus, we tested whether *legK2* and *vipA* genes have evolved independently of each other or co-evolved in the genus *Legionella*. We examined the occurrences of LegK2 and VipA across 46 *Legionella* species (and *Coxiella burnetti* as an outgroup) represented by a set of 647 genomes characterized by an over-representation of *L. pneumophila* genomes (540/647) **(Supplementary Data S9)**. Strikingly, the LegK2/VipA pair was restricted to *L. pneumophila* species, and VipA was only present in a clade closely related to *L. pneumophila*, containing *L. waltersii, L. moravica, L. quateirensis, L. shakespearei and L. worsleiensis* **(figure 5A)**. The absence of the LegK2/VipA pair in the rest of the genus *Legionella* is consistent with a previous report that showed that only 7 Dot/Icm effectors were conserved in 41 *Legionella* species, confirming the high versatility of the effector repertoire^39^. More interestingly, in *L. pneumophila* the co-occurrence of LegK2 and VipA is highly frequent (in 96.48 % of the *L. pneumophila* genomes), as, among the 540 genomes of *L. pneumophila* included in our study, only 7 possess LegK2 only and 12, VipA only **(figure 5B)**. We investigated whether strains that had lost either LegK2 or VipA differed from other *L. pneumophila* strains in their conservation of other effectors known to interfere with the actin cytoskeleton. Phylogenetic study of LegK2, VipA, Ceg14, RavK and WipA reveals that this set of effectors is conserved in *L. pneumophila* species as the five effectors are found together in 512 of the 540 *L. pneumophila* genomes (94%) **(Supplementary Data S10)** and Ceg14, RavK and WipA are conserved in the 18 strains that have lost LegK2 or VipA **(Supplementary Data S10)**. This complete set of effectors is most likely the result of an evolutionary history that selected the effector repertoire best suited to manipulate host actin polymerization to the benefit of the bacterium. Finally, we used Count^28^ to reconstruct the evolutionary history of the LegK2/VipA pair, onto a tree of 120 genomes representing the phylogenetic diversity of *Legionella* sp. while biasing our sample towards the 19 *L. pneumophila* genomes missing one of each gene. The analysis confirmed that VipA and LegK2 are both ancestral to *L. pneumophila*, but also revealed that VipA was acquired before LegK2. VipA originated in the ancestor of a clade grouping *L. pneumophila, L. waltersii, L. moravica, L. quateirensis, L. shakespearei and L. worsleiensis*, while LegK2 was specifically acquired in the ancestor of *L. pneumophila* species. Subsequently, either LegK2 or VipA were sporadically and independently lost **(figure 5A)**.

**Figure 5.**
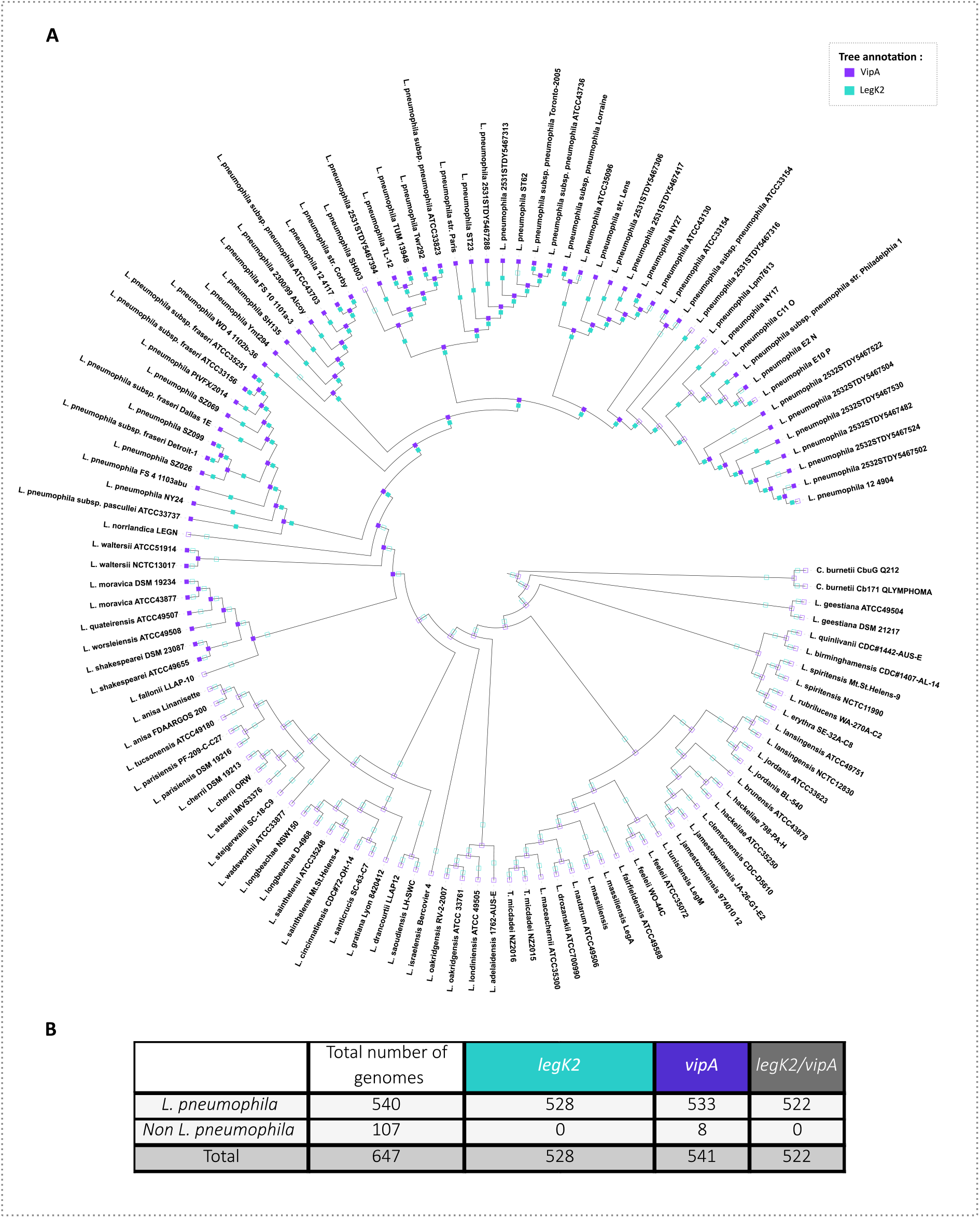
LegK2/VipA effector pairs are restricted and strongly conserved in the *L. pneumophila* species. **(A)** Phylogenetic tree representing the phylogenetic diversity of *Legionella* genomes displaying either LegK2 (blue green squares) or VipA genes (purple squares). The squares are filled when the genes have been found in corresponding genome and empty when genes have not been detected. **(B)** Distribution of 2 selected effectors from *Legionella pneumophila* in 647 different *Legionella* strains including *L. pneumophila* and non *pneumophila* strains. The protein sequence of the effector of *L. pneumophila* strain was blasted against our genome database of family-clustered proteins to determine the presence or absence of proteins in the corresponding genomes.

## DISCUSSION

*L. pneumophila* has evolved the largest arsenal of bacterial effectors to control a sophisticated relationship with its host phagocytic cells, amoebae or human macrophages. The record number of over 300 effectors was hypothesized to result from co-evolution with its highly diverse environmental hosts to set up the best effector repertoire for each of its host. It is assumed that many effectors are needed to orchestrate complex and sequential interactions with numerous host cell pathways in order to support the intracellular multiplication of the bacterium. Recently, systematic screens^19,40^ or individual studies of effector function^41–49^ have identified some effectors that interfere with other effector activity rather than targeting a host cell protein. Regardless the specific activities involved in these effector-effector interactions, two main models of interaction were revealed: indirect through counteracting modification of a shared host target, such as an effector pair recently described as para-effectors that target the host cell histone H3^50^; or direct through either complex formation or the modification of one effector by another. The direct interaction model led to the emerging concept of « metaeffector ».

Our study contributes to this field by identifying a novel effector-effector suppression pair, namely LegK2/VipA, which targets host actin cytoskeleton to support bacterial evasion from endosomal degradation and subsequent intracellular replication. This effector-effector suppression pair was not revealed by the high-throughput systematic screen based on the rescue of yeast growth defect upon heterologous expression of *L. pneumophila* effectors^19^, most likely because ectopic expression of either LegK2 or VipA (yet identified to disrupt membrane trafficking in yeast^51^), does not alter yeast growth sufficiently to be detectable in the screen. Importantly, the LegK2/VipA suppression pair is physiologically relevant since it has been here identified in physiological conditions of infection of both amoeba and macrophages. Specifically, we demonstrated that *vipA* gene deletion rescues the *ΔlegK2* defects of inhibition of phagosome maturation along the endosomal pathway and bacterial intracellular multiplication. LegK2 and VipA antagonistic activities towards actin polymerization, i.e. inhibition of actin nucleation by LegK2 through the targeting of Arp2/3 complex *versus* direct actin nucleation by VipA, result in controlling actin polymerization on the LCV. In addition to the identification of the novel LegK2/VipA suppression pair, our data propose for the first time a role for VipA in the infectious cycle of *L. pneumophila*; thus, identification of effector-effector suppression pairs could be effective in circumventing the significant functional redundancy of effectors and in gaining insights into the unknown function of many Dot/Icm effectors.

The functional interaction between LegK2 and VipA is consolidated by the evolutionary biology of this pair of effectors. Indeed, both effectors are restricted to *L. pneumophila* species and are found together at a frequency of 96.48 % in the different strains of *L. pneumophila*. VipA was acquired by the clade containing *L. pneumophila*, but also *L. waltersii, L. moravica, L. quateirensis, L. shakespearei* and *L. worsleiensis* before LegK2 was specifically acquired by the *L. pneumophila* species. Then, some independent events of loss of each gene occurred very sporadically among the *L. pneumophila* phylogeny. Noteworthy, the reductive model of laboratory infection assays involving a very small number of amoeba species (usually *A. castellanii*), thereby reducing the environmental host diversity of *L. pneumophila*, does not allow for an assessment of the impact of these loss events on the intracellular replication of these *L. pneumophila* strains. Overall, the evolutionary history of LegK2/VipA pair excludes the initial model of acquisition of effector-effector suppression pairs through a common horizontal gene transfer. It is consistent with the distant localization of *legK2* and *vipA* genes on the *L. pneumophila* genome, and likely similar for some other effector-effector suppression pairs encoded by distant genes ^19^.

LegK2 and VipA are likely expressed and secreted at the same time into the host cell, specifically in the transmissive phase of the *L. pneumophila* infection cycle, and they would be available to interfere with the host cell pathway at the early stage of the infection. As Urbanus *et al*. suggest, transcriptomics is not sufficient to place effector-effector interaction in the context of infection, and detailed proteomic analyses and secretion assays that would reveal potential controls of translation, secretion and localization of the effectors in the cell are the next step in the field ^19^. Indeed, while the *vipA* gene is over-expressed in the exponential phase, the VipA protein appears to accumulate in the bacterium in the post-exponential growth phase. Despite their patterns of co-expression and co-secretion, the LegK2/VipA suppression pair does not meet the definition of metaeffector. They do not seem to physically interact, or colocalize inside the eukaryotic cells. Moreover, despite the fact that some metaeffectors have already been shown to target both host cell proteins and other bacterial effectors ^19,52^, VipA is not a phosphorylation substrate of the protein kinase LegK2. Thus, our data on the LegK2/VipA suppression pair provide an additional model of effector-effector suppression interaction in which two effectors with antagonistic activities towards distinct actors of a same host cell pathway finely cooperate to modulate the host cell to the benefit of the bacterium. Given that LegK2 is detected on the LCV surface 1 h post-infection ^16^ and that VipA-derived peptides were identified at the LCV 1 h post-infection ^38^, it can be hypothesized that the antagonistic activities of the LegK2/VipA effector pair can temporally control actin polymerization on the LCV to interfere with phagosome maturation and endosome recycling. Indeed, the actin polymerization inhibition activity exerted by LegK2, followed by the actin polymerization activity exerted by VipA, would be consistent with the well-established observation that, after bacterial engulfment, actin and associated proteins dissembles but are recruited again at a later stage of endocytic transit ^30,53,54^. In addition to directly controlling actin polymerization, *L. pneumophila* also controls actin degradation at the LCV ^55^ and secretes RavK, an actin-targeting effector protease. To achieve a comprehensive model of host actin cytoskeleton manipulation by *L. pneumophila*, the full set of 5 effectors known to interfere with actin polymerization (LegK2, VipA, Ceg14, RavK and WipA) must be considered, and functional interactions between them will be studied in more detail.

## CONCLUSION

The delivery of effector proteins that hijack host cell processes for the benefit of the bacteria is a mechanism widely used by bacterial pathogens and thus plays a key role in microbial virulence. *L. pneumophila*, which translocates a record number of 300 effectors, is the paradigm of a pathogen that have evolved a highly sophisticated relationship with its hosts, and a perfect model to study the complex action of effectors and their functional interactions. Here, we revealed a new type of effector-effector suppression pair in *L. pneumophila*, which neither meets the emerging definition of a metaeffector, i.e. an effector that controls the activity of another effector, nor the new definition of a paraeffector, i.e. two effectors that act synergistically or oppositely on the same cellular target. Instead, LegK2 and VipA express their actin polymerization antagonistic activities by targeting two distinct cellular targets through two different molecular mechanisms, in order to finely control actin polymerization on the LCV surface. Importantly, their combined actions play a key role in the escape of bacteria from endocytic degradation and subsequent intracellular bacterial replication.

## Acknowledgments

We thank Adeline Page and Frédéric Delolme from the Protein Science Facility at the SFR Biosciences (UAR3444/CNRS, US8/Inserm, ENS de Lyon, UCBL) for the Mass spectrometry analyses. We are grateful to F. Letourneur for the gift of anti-VatA antibody. We thank the Dicty Stock Center for *D. discoideum* strains. And finally, we thank Camille Fourneaux (LBMC, ENS de Lyon) for helping with R scripts.

## Author contributions

P.D. designed research; M.P., C.M., and N.B. performed experimental research, except proteomics; L.A. and K.P. designed, performed and analysed proteomics; M.P., J.B., E.K., and V.D. designed, performed and analyse evolutionary biology. M.P., C.M., N.B., O.D. and P.D. analysed data; M.P. and P.D. wrote the paper.

## Data availability statement

RNA-seq and sequencing data have been deposited in European Nucleotide Archive database at EMBL-EBI (https://www.ebi.ac.uk/ena) under accession number PRJEB62121. The data that support the findings of this study are available from the corresponding author upon reasonable request.

