## Supplementary Data for "Dual control of host actin polymerization by a *Legionella* effector pair"

### SUPPLEMENTAL DATA LEGENDS

**Supplementary Data S1 – List of all cells, bacteria strains used in this article.**

**Supplementary Data S2 – List of plasmids used in this study.**

**Supplementary Data S3 – List of oligonucleotides used to realize mutant strains of *Legionella* as well as for cloning.**

**Supplementary Data S4 – Accession numbers of genomes used for evolutionary analyses.**

**Supplementary Data S5 – Fasta files of sequencing data of *ΔlegK2* and *ΔlegK2/ΔvipA Legionella pneumophila* strains.** RNA-seq and sequencing data have been deposited in European Nucleotide Archive database at EMBL-EBI (https://www.ebi.ac.uk/ena) under accession number PRJEB62121.

**Supplementary Data S6 – LegK2 and VipA mutations do not alter axenic growth of *L. pneumophila* Paris strains.** Axenic growth kinetics of L. pneumophila Paris WT, Δ*dotA,* Δ*legK2,* Δ*vipA,* Δ*legK2/*Δ*vipA, legK2/vipA*ΔNter and *legK2/vipA*ΔCter mutant *L. pneumophila* strains transformed with mCherry-expressing plasmids. AYE broth was seeded with 0,1 $x10^9$ bacteria/mL and then plated in a 96well-Greiner plate, with black wells and transparent flat bottom. $OD_{600nm}$ absorbance was measured every 30 min at 595 nm on an Infinite M200 microplate reader (Tecan).

**Supplementary Data S7 – Mass Spectrometry data following protein expression in *Legionella pneumophila* Paris WT at $OD_{600nm}$ 1, 2, 3, 4 and 5.** VipA accumulates in *L. pneumophila* up to $DO_{600}$= 4. Three independent cultures of *L. pneumophila* Paris WT were grown in liquid medium AYE (at 30°C). At the desired $OD_{600}$ (1, 2, 3, 4 and 5), samples were collected, and their protein content was analysed by mass spectrometry after sample-specific labelling (see Material and methods for details). The graphs show the quantity detected at each $OD_{600}$ or as a ratio compared to the quantity at $OD_{600}$ 3. The RocC protein is used as a control as it is detected at the same range of quantity as VipA and its production during growth was previously studied by Western-blot [23]. Of note the RocC pattern of production detected by mass spectrometry corresponds to the one previously obtained by Western blot[23].

**Supplementary Data S8 – Localization during infection of HA-LeK2 and HA-VipA in *D. discoideum*. (A and B)** *D. discoideum* were infected at MOI = 50 for 1h30 with *L. pneumophila* WT Paris transformed with pMMB207c-Ptac-HA-*legK2* or pMMB207c-Ptac-HA-*vipA* plasmid. At each time point, infected cells were fixed, and immunofluorescence was carried out as described (Materials and Methods). DNA was stained with DAPI, HA-LegK2 **(A)**/HA-VipA **(B)** with an anti-HA antibody. Scale = 10 µm. **(C)** Control experiment was performed, infecting *D. discoideum* at MOI = 50 for 1 hour using *L. pneumophila* WT Paris plasmid or secretion-deficient mutant Δ*dotA* transformed with pMMB207c-Ptac-HA-*legK2* or pMMB207c-Ptac-HA-*vipA* as a control for secretion. As expected, no fluorescence was detected for either condition which demonstrates that the procedure does not permeabilize bacteria. **(D)** *D. discoideum* were infected at MOI = 50 for 24h with *L. pneumophila* WT Paris transformed with pMMB207c-Ptac-HA-*vipA* plasmid. At 24h post-infection, infected cells were fixed, and immunofluorescence was carried out as described (Materials and Methods). DNA was stained with DAPI, HA-VipA with an anti-HA antibody. Scale = 5 µm. HA-VipA was detected in higher amount in the infected cells cytosol at 24h post-infection.

**Supplementary Data S9 – Coomassie blue staining gels to check the overexpression and purification of GST-LegK2 and 6His-VipA.** GST-LegK2 and 6His-VipA were purified as described in Material and Methods. *E. coli* BL21 (DE3) strains carrying either pGEX-legK2 or pQE30-VipA were grown overnight and backdiluted 1:100 and cultivated at 37°C until cultures reached an $OD_{600\ nm}$ of 0.7. Then, IPTG was added to a final concentration of 0,2 mM and growth was continued overnight at 20°C. The next day, pellets were resuspended in GST-Pull Down equilibration/wash buffer (125 mM Tris-HCl pH 7.5, 150 mM NaCl + Protease Inhibitor Cocktail (Sigma) + 1 mg/ml lysozyme) or in 6His-Pull Down equilibration/wash buffer (50 mM Tris-HCl pH 7.5, 150 mM NaCl and 10 mM Imidazole + Protease

Inhibitor Cocktail (Sigma) + 1 mg/ml lysozyme) and bacteria were lysed using 3 passages in a French Press (SLM, Urbana, IL) and lysates centrifugated at 12 000 rpm during 30 min. Supernatants were then collected and transferred to tubes containing either Pierce™ Gluthatione Magnetic Agarose Beads (ThermoFisher Scientific) or TALON Metal Affinity Resin (Takara Bio) respectively, according to the manufacturers' recommendations. The purity of the eluted protein was analysed by SDS-PAGE.

**Supplementary Data S10 – Other effectors targeting actin polymerization, Ceg14, WipA and RavK are strongly conserved in the *L. pneumophila* species. (A)** Distribution of the 5 actin-polymerization related- effectors from *Legionella pneumophila* in 647 different *Legionella* strains including *L. pneumophila* and non *pneumophila* strains. The protein sequence of the effector of *L. pneumophila* strain was blasted against our genome database of family-clustered proteins to determine the presence or absence of proteins in the corresponding genomes. **(B)** Phylogenetic tree representing the phylogenetic diversity of *Legionella* genomes displaying either LegK2 (name of species written in blue), VipA (written in purple), Ceg14 (green squares), RavK (yellow squares) and WipA genes (dark blue). The squares are filled when the genes have been found in corresponding genome and empty when genes have not been detected.

**SUPPLEMENTARY DATA**

| **Strains** | | |
|---|---|---|
| **Names** | **Genotypes** | **References** |
| ***Dictyostelium discoideum*** | | |
| DBS0235534 | Ax2-214 | (Watts & Ashworth, 1970) |
| ***Acanthamoebae castellanii*** | | |
| Environmental isolate | | |
| **Mammalian cells** | | |
| HeLa cells | | Inserm U1111, Lyon, France |
| U937 | Human monocytes ATCC CRL1593.2 | Sundstrom et al. 1976 |
| ***Legionella pneumophila*** | | |
| WT | Virulent *L. pneumophila* strain Paris CIP 107629 | Cazalet et al. (2004) |
| *ΔdotA* | Paris *lpp2740*::Km | Cazalet et al. (2004) |
| *ΔlegK2* | Paris Δ*lpp2076* (scar-free deletion of lpp2076) | This study |
| *ΔvipA* | Paris Δ*lpp0457* (scar-free deletion of lpp0457) | This study |
| *ΔlegK2ΔvipA* | Paris Δ*lpp2076* Δlpp0457 (scar-free deletion of lpp2076 and lpp0457) | This study |
| ***Escherichia coli*** | | |
| DH5α | *endA1 hsdR17 supE4, thi-1 recA1 gyrA relA, Δlac* | Laboratory |
| BL21 | B *dcm ompT hsdS(rB-mB-) gal* | Laboratory |
| XL1 Blue | *endA1 gyrA96(nalR) thi-1 recA1 relA1 lac glnV44* F'[/Tn10 *proAB* + *lacIq*Δ*(lacZ*)M15] *hsdR17*(rK-mK+) | Stratagene |

**SUPPLEMENTARY DATA S1**

| Plasmids | | |
|---|---|---|
| **Names** | **Characteristics** | **References** |
| Donor vectors for Gateway cloning | | |
| pDONR™207 | Donor Gateway vector | Invitrogen |
| pDONR™207-*vipA* | Donor Gateway vector with insertion of lpp0457 gene | This study |
| Mammalian cells expression vectors for Gateway cloning | | |
| pDEST27 | Expression Gateway vector for mammalian cells allowing overexpression of GST-tagged protein | Invitrogen |
| pDEST27-*legK2* | pDEST27 with insertion of lpp2076 gene for expression of GST-LegK2 fusion protein | This study |
| pDEST27-*legK2*$_{K112M}$ | pDEST27 with insertion of lpp2076 gene for expression of GST-LegK2$_{K112M}$ dead kinase-mutant fusion protein | This study |
| pCI-Neo3Flag | Expression Gateway vector for mammalian cells allowing overexpression of Flag-tagged protein | Invitrogen |
| pCI-Neo3Flag-*ArpC1b* | pCI-Neo3Flag with insertion of ARPC1B human gene for expression of Flag-ARPC1B fusion protein | (Michard et al. 2015) |
| peGFP-Nterm | Expression Gateway vector for mammalian cells allowing overexpression of Nterminal GFP-tagged protein | INSERM U1111, Lyon, France |
| peGFP-Cterm | Expression Gateway vector for mammalian cells allowing overexpression of Nterminal GFP-tagged protein | INSERM U1111, Lyon, France |
| peGFP-N-*vipA* | peGFP-Nterm with insertion of lpp0457 gene for expression of VipA-GFP fusion protein | This study |
| peGFP-C-*vipA* | peGFP-Cterm with insertion of lpp0457 gene for expression of GFP-VipA fusion protein | This study |
| Bacteria expression vectors | | |
| pXDC50 | Expression plasmid for *L. pneumophila* allowing expression of mCherry fluorescent protein | (Hervet et al. 2011) |
| pXDC61 | Expression plasmid for *L. pneumophila* allowing overexpression of β-Lactamase | |
| pXDC61-*legK2* | Expression plasmid for *L. pneumophila* allowing overexpression of β-Lactamase-LegK2 fusion protein | This study |
| pXDC61-*vipA* | Expression plasmid for *L. pneumophila* allowing overexpression of β-Lactamase-VipA fusion protein | This study |
| pGEX-*legK2* | Overproduction and *in vitro* purification of GST-LegK2 | This study |
| pQE30-*vipA* | Overproduction and *in vitro* purification of GST-LegK2 | This study |

**SUPPLEMENTARY DATA S2**

| Primers | | |
|---|---|---|
| **Names** | **Sequences** | **Description** |
| 1_vipA fwd | GGGGACAAGTTTGTACAAAAAAGCAGGCTTAatgcctatcagtaatgccttt | *vipA* insertion in pDONR207 vector for peGFP-C-VipA |
| 2_vipA rev | GGGGACCACTTTGTACAAGAAAGCTGGGTTgagattttttttttcgacggtagtg | |
| 3_vipA fwd | GGGGACCACTTTGTACAAGAAAGCTGGGTTgagattttttttttcgacggtagtg | *vipA* insertion in pDONR207 vector for peGFP-N-VipA |
| 4_vipA rev | GGGGACCACTTTGTACAAGAAAGCTGGGTTgagattttttttttcgacggtagtg | |
| 5_legK2 fwd | attggggaagcggtacccgggtttattacataaatttgaagga | *legK2* insertion in pXDC61 vector in XmaI site |
| 6_legK2 rev | tatctagaggatccccgggttttagcttgggcctcgcat | |
| 7_vipA fwd | cattggggaagcggtacccggcctatcagtaatgcctttcttaa | *vipA* insertion in pXDC61 vector in XmaI site |
| 8_vipA rev | tatctagaggatccccggctagagattttttttttcgacggta | |
| 9_P1 vipA | TTCAACTCATCCTAGCATTG | *vipA* inactivation in *Legionella* by KanMazF technic |
| 10_P2 vipA | CAGCAATATGATATTCTGGC | |
| 11_P3 vipA | GGCCCAATTCGCCCTATAGTGAGTCGATGGATGGTCAAGCATTATC | |
| 12_P4 vipA | GGGTTTGCTCGGGTCGGTGGCATATGGTGCTCTATGATACTGACATC | |
| 13_P5 vipA | GATGTCAGTACATAGAGCACATGGATGGTCAAGCATTATC | |
| 14_P6 vipA | GATAATGCTTGACCATCCATGTGCTCTATGATACTGACATC | |
| 19_P1 legK2 | TTACCAATGTAATGAGACATCG | *legK2* inactivation in *Legionella* by KanMazF technic |
| 20_P2 legK2 | GGCCCAATTCGCCCTATAGTGAGTCGtcttgaggtagaggttgttcc | |
| 21_P3 legK2 | GGGTTTGCTCGGGTCGGTGGCATATGCTGCAAATCAAGATAAGCAACC | |
| 22_P4 legK2 | ATAGCATGCACAACTTTGATATCAAGCATCC | |
| 23_P5 legK2 | GGTTGCTTATCTTGATTTGCAGtcttgaggtagaggttgttcc | |
| 24_P6 legK2 | GGAACAACCTCTACCTCAAGACTGCAAATCAAGATAAGCAACC | |
| 25_Fwd pGEX | GGGCCCCTGGAACAGAAC | Construction of pGEX-*legK2* plasmid by PCR SLIC |
| 26_Rev pGEX | CTGGGATCCCCGAATTCCC | |
| 27_Fwd legK2 pGEX | GTTCTGTTCCAGGGGCCCgtttattacataaatttgaaggaac | |
| 28_Rev legK2 pGEX | GGGAATTCGGGGATCCCAGttagcttgggcctcgcatc | |

| | | |
|---|---|---|
| 31_Fwd vipA pQE30 | GCTCGCATGCGGATCctagagattttttttttcgacggtag | Construction of pQE30-*vipA* plasmid by BamHI digestion SLIC |
| 32_Rev vipA pQE30 | TCACCATCACGGATCccctatcagtaatgcctttcttaagtt | |

**SUPPLEMENTARY DATA S3**

| Accession # | Strain names | Accession # | Strain names |
|---|---|---|---|
| **GCF000621365.1** | *L. geestiana* DSM 21217 | **GCF001468085.1** | *L. waltersii* ATCC51914 |
| **GCF001467645.1** | *L. geestiana* ATCC49504 | **GCF900187095.1** | *L. waltersii* NCTC13017 |
| **GCF001468165.1** | *L. spiritensis* Mt.St.Helens-9 | **GCF000770585.1** | *L. norrlandica* LEGN |
| **GCF900186965.1** | *L. spiritensis* NCTC11990 | **GCF001582625.1** | L. *pneumophila* FS_4_1103abu |
| **GCF001467615.1** | *L. erythra* SE-32A-C8 | **GCF000586195.1** | L. *pneumophila* subsp. fraseri ATCC35251 |
| **GCF001468125.1** | *L. rubrilucens* WA-270A-C2 | **GCF000586315.1** | L. *pneumophila* subsp. fraseri ATCC33156 |
| **GCF001467505.1** | *L. birminghamensis* CDC#1407-AL-14 | **GCF001639045.1** | *L. pneumophila* PtVFX/2014 |
| **GCF001467975.1** | *L. quinlivanii* CDC#1442-AUS-E | **GCF001582535.1** | *L. pneumophila* SZ069 |
| **GCF001467025.1** | *L. brunensis* ATCC43878 | **GCF001886835.1** | *L. pneumophila* subsp. fraseri Dallas 1E |
| **GCF000953655.1** | *L. hackeliae* ATCC35250 | **GCF001582295.1** | *L. pneumophila* SZ099 |
| **GCF001467705.1** | *L. hackeliae* 798-PA-H | **GCF001886795.1** | *L. pneumophila* subsp. fraseri Detroit-1 |
| **GCF002240035.1** | *L. clemsonensis* CDC-D5610 | **GCF001583645.1** | *L. pneumophila* SZ026 |
| **GCF001691475.1** | *L. jamestowniensis* 974010_12 | **GCF001600905.1** | *L. pneumophila* NY24 |
| **GCF001467745.1** | *L. jamestowniensis* JA-26-G1-E2 | **GCF000586255.1** | L. pneumophila *subsp. pascullei* ATCC33737 |
| **GCF900187355.1** | *L. lansingensis* NCTC12830 | **GCF001582645.1** | *L. pneumophila* WD_4_1102b-36 |
| **GCF001467795.1** | *L. lansingensis* ATCC49751 | **GCF000953935.1** | *L. pneumophila* Ymt294 |
| **GCF001467765.1** | *L. jordanis* BL-540 | **GCF001582405.1** | *L. pneumophila* FS_10_1101a-3 |
| **GCF001648675.1** | *L. jordanis* ATCC33623 | **GCF000092625.1** | *L. pneumophila* 2300/99 Alcoy |
| **GCF000308315.1** | *L. tunisiensis* LegM | **GCF000092545.1** | *L. pneumophila* str. Corby |
| **GCF001467625.1** | *L. feeleii* WO-44C | **GCF000823425.1** | *L. pneumophila* 12_4117 |
| **GCF001648615.1** | *L. feeleii* ATCC35072 | **GCF000586095.1** | *L. pneumophila subsp. pneumophila* ATCC43703 |
| **GCF000621525.1** | *L. fairfieldensis* ATCC49588 | **GCF001582385.1** | *L. pneumophila* SH135 |
| **GCF000756695.1** | *L. massiliensis* | **GCF900063795.1** | *L. pneumophila* 2531STDY5467313 |
| **GCF000756815.1** | *L. massiliensis* LegA | **GCF000586075.1** | *L. pneumophila subsp. pneumophila* ATCC43736 |
| **GCF001467585.1** | *L. drozanskii* ATCC700990 | **GCF001592705.1** | *L. pneumophila subsp. pneumophila* Toronto-2005 |
| **GCF001467895.1** | *L. nautarum* ATCC49506 | **GCF900119765.1** | *L. pneumophila* ST62 |
| **GCF900167045.1** | *L. maceachernii* ATCC35300 | **GCF900053665.1** | *L. pneumophila* 2531STDY5467288 |
| **GCF002085735.1** | *T. micdadei* NZ2016 | **GCF900119755.1** | *L. pneumophila* ST23 |
| **GCF002085715.1** | *T. micdadei* NZ2015 | **GCF000048645.1** | *L. pneumophila* str. Paris |
| **GCF001467055.1** | *L. adelaidensis* 1762-AUS-E | **GCF001583655.1** | *L. pneumophila* SH003 |
| **GCF001467825.1** | *L. londiniensis* ATCC 49505 | **GCF900060375.1** | *L. pneumophila* 2531STDY5467394 |
| **GCF000512715.1** | *L. oakridgensis* RV-2-2007 | **GCF000695015.1** | *L. pneumophila* TUM 13948 |
| **GCF001648605.1** | *L. oakridgensis* ATCC 33761 | **GCF001582555.1** | *L. pneumophila* TL-12 |
| **GCF001467785.1** | *L. israelensis* Bercovier 4 | **GCF000586235.1** | *L. pneumophila subsp. pneumophila* ATCC33823 |
| **GCF000162755.2** | *L. drancourtii* LLAP12 | **GCF000950745.1** | *L. pneumophila* Twr292 |
| **GCF001465875.1** | *L. saoudiensis* LH-SWC | **GCF001582225.1** | *L. pneumophila* ATCC35096 |
| **GCF001467695.1** | *L. gratiana* Lyon 8420412 | **GCF000306865.1** | *L. pneumophila subsp. pneumophila* Lorraine |
| **GCF001467545.1** | *L. cincinnatiensis* CDC#72-OH-14 | **GCF900050185.1** | *L. pneumophila* 2531STDY5467306 |
| **GCF001468135.1** | *L. santicrucis* SC-63-C7 | **GCF001582215.1** | *L. pneumophila* ATCC43130 |

| | | | |
|---|---|---|---|
| **GCF000621685.1** | *L. sainthelensi* ATCC35248 | **GCF001601075.1** | *L. pneumophila* NY27 |
| **GCF001468105.1** | *L. sainthelensi* Mt.St.Helens-4 | **GCF900062515.1** | *L. pneumophila* 2531STDY5467417 |
| **GCF000091785.1** | *L. longbeachae* NSW150 | **GCF000048665.1** | *L. pneumophila* str. Lens |
| **GCF000176095.1** | *L. longbeachae* D-4968 | **GCF000586355.1** | *L. pneumophila subsp. pneumophila* ATCC33154 |
| **GCF000701265.1** | *L. wadsworthii* ATCC33877 | **GCF001582565.1** | *L. pneumophila* ATCC33154 |
| **GCF001467945.1** | *L. parisiensis* PF-209-C-C2 | **GCF900057235.1** | *L. pneumophila* 2531STDY5467316 |
| **GCF001736145.1** | *L. parisiensis* DSM 19216 | **GCF900092465.1** | *L. pneumophila* Lpm7613 |
| **GCF001468035.1** | *L. tucsonensis* ATCC49180 | **GCF001753125.1** | *L. pneumophila* C11_O |
| **GCF000333755.1** | *L. anisa* Linanisette | **GCF001752705.1** | *L. pneumophila* E10_P |
| **GCF002082905.1** | *L. anisa* FDAARGOS_200 | **GCF001753265.1** | *L. pneumophila* E2_N |
| **GCF001468065.1** | *L. steigerwaltii* SC-18-C9 | **GCF000008485.1** | *L. pneumophila subsp. pneumophila* str. Philadelphia 1 |
| **GCF000621385.1** | *L. cherrii* DSM 19213 | **GCF001601055.1** | *L. pneumophila* NY17 |
| **GCF001467035.1** | *L. cherrii* ORW | **GCF900058585.1** | *L. pneumophila* 2532STDY5467522 |
| **GCF001468005.1** | *L. steelei* IMVS3376 | **GCF900062435.1** | *L. pneumophila* 2532STDY5467504 |
| **GCF000953135.1** | *L. fallonii* LLAP-10 | **GCF900062395.1** | *L. pneumophila* 2532STDY5467530 |
| **GCF000373765.1** | *L. shakespearei* DSM 23087 | **GCF000823645.1** | *L. pneumophila* 12_4904 |
| **GCF001468025.1** | *L. shakespearei* ATCC49655 | **GCF900055205.1** | *L. pneumophila* 2532STDY5467502 |
| **GCF001467535.1** | *L. worsleiensis* ATCC49508 | **GCF900062385.1** | *L. pneumophila* 2532STDY5467524 |
| **GCF000423305.1** | *L. moravica* DSM 19234 | **GCF900053395.1** | *L. pneumophila* 2532STDY5467482 |
| **GCF001467865.1** | *L. moravica* ATCC43877 | **GCF000826165.1** | *C. burnetii* Cb171_QLYMPHOMA |
| **GCF001467955.1** | *L. quateirensis* ATCC49507 | **GCF000019865.1** | *C. burnetii* CbuG_Q212 |

### SUPPLEMENTARY DATA S4

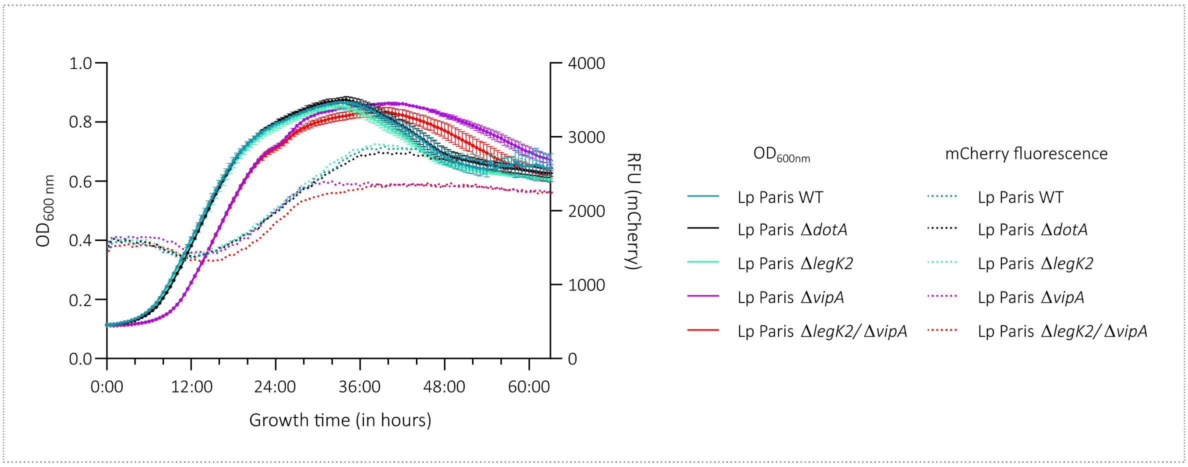


**SUPPLEMENTARY DATA S**

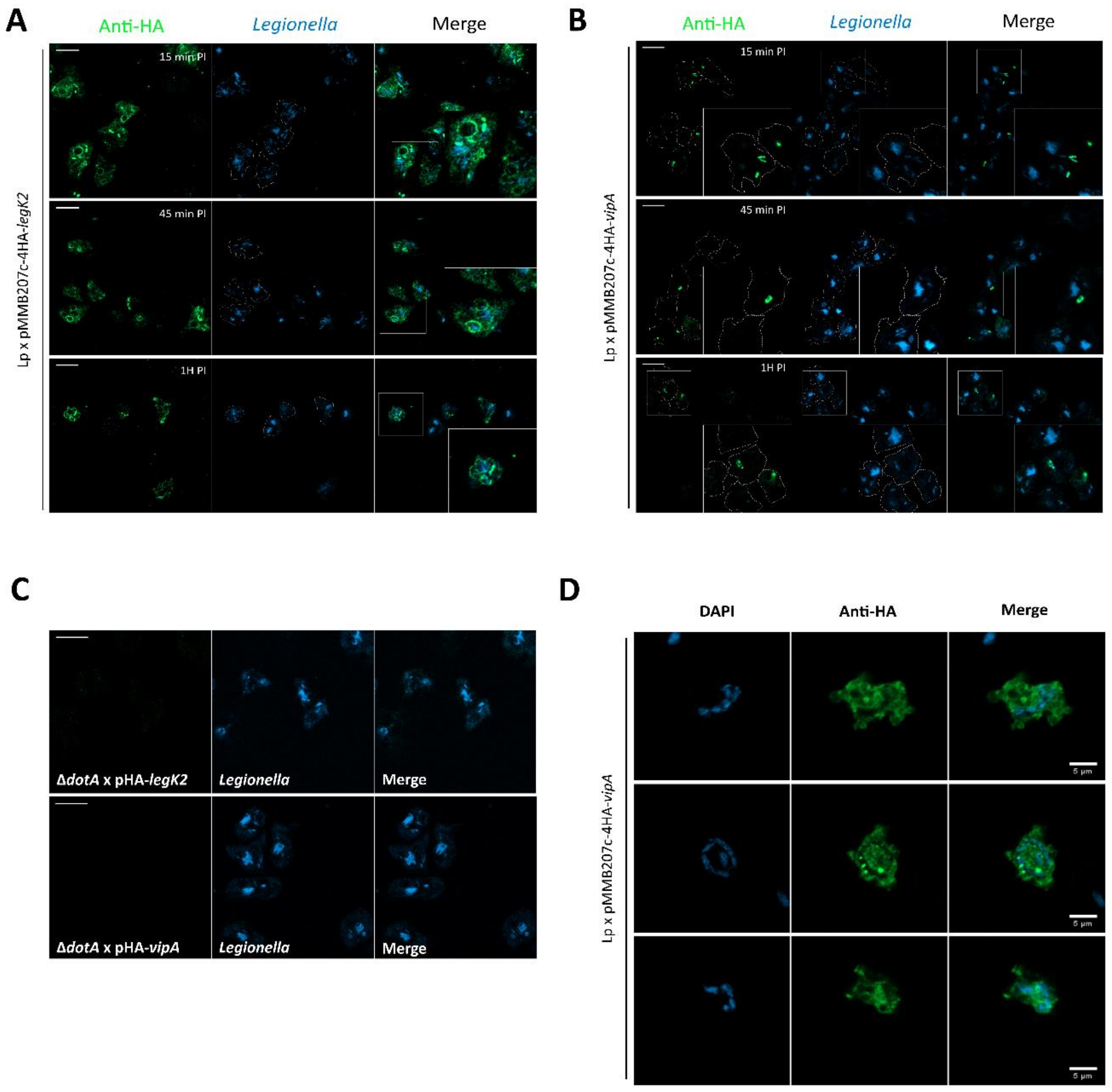


**SUPPLEMENTARY DATA S8**

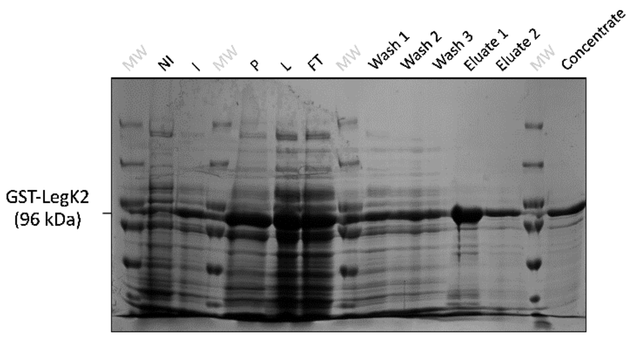


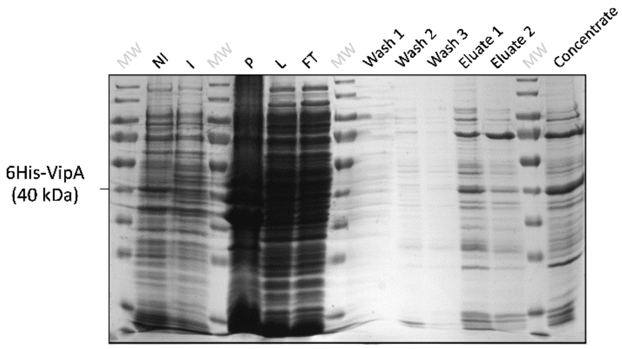


**SUPPLEMENTARY DATA S9**

**A**

| Co-occuring genes | *L. pneumophila* | *Non L. pneumophila* |
|---|---|---|
| Total number of genomes | 540 | 107 |
| **All 5** | 512 | 0 |

**B**

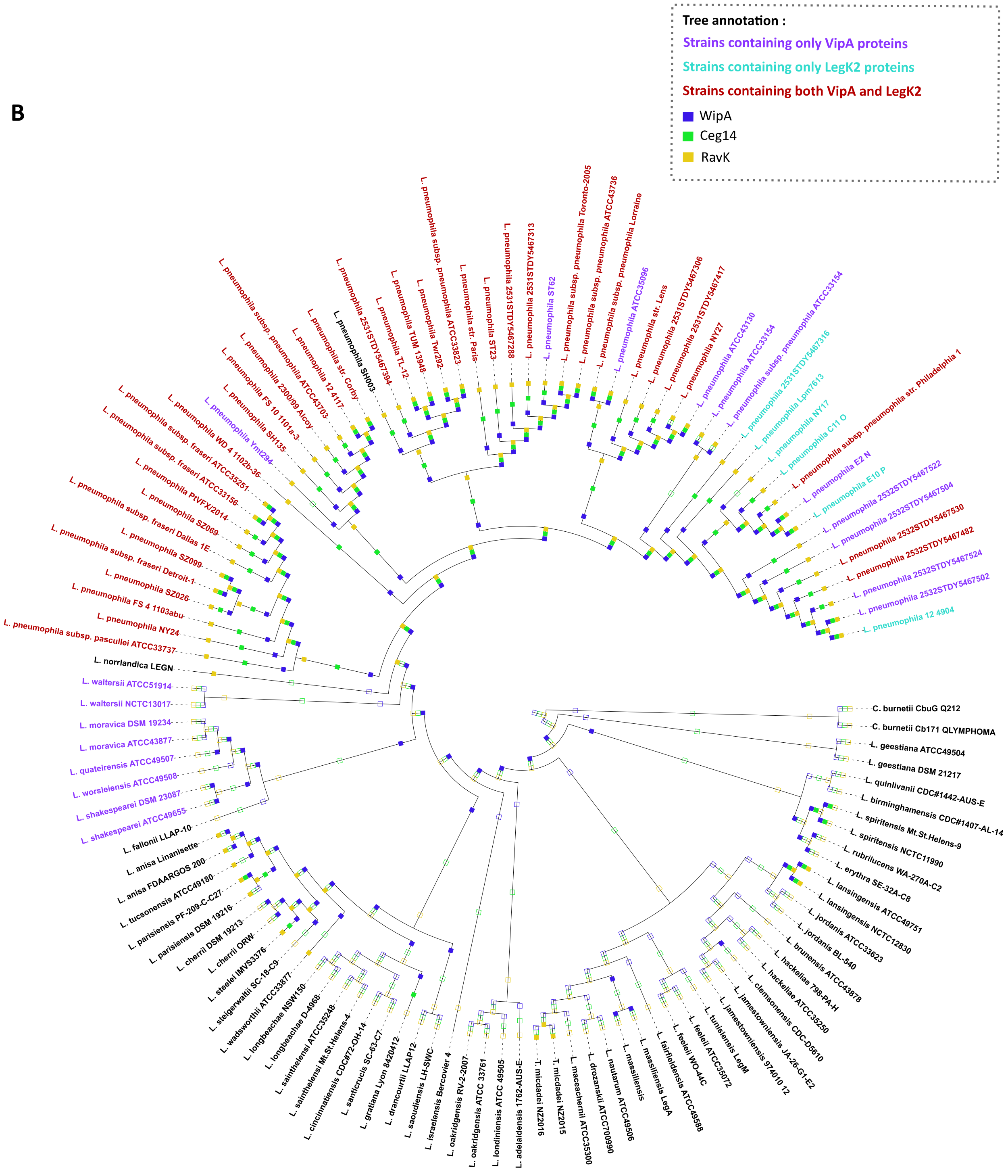


### SUPPLEMENTARY DATA S10
